# Expression of a modified astrocytic glutamate transporter alleviates Huntington’s hypokinesia, promotes synaptic glutamate clearance and counteracts potentially adverse EAAT2 interactions

**DOI:** 10.1101/2020.09.17.302158

**Authors:** Stefan Hirschberg, Anton Dvorzhak, Seyed M. A. Rasooli-Nejad, Svilen Angelov, Marieluise Kirchner, Philip Mertins, Gilla Lättig-Tünnemann, Christoph Harms, Dietmar Schmitz, Rosemarie Grantyn

## Abstract

Rapid removal of glutamate from the sites of glutamate release is an essential step in excitatory synaptic transmission. Despite many years of research, the molecular mechanisms underlying the intracellular regulation of glutamate transport at tripartite synapses have remained unclear. This limits the options for pharmacological treatment of motor disorders associated with glutamate excitotoxicity. Therefore, using the Q175 mouse model of Huntington’s disease (HD), we explored the effects of structural changes in the astrocytic excitatory amino acid transporter type 2 (EAAT2). We report that expression of a C-terminal-modified variant of EAAT2 can alleviate the symptoms of hypokinesia in mice with already advanced HD. At a cellular level, this beneficial outcome correlated with faster synaptic glutamate clearance, higher astrocytic glutamate uptake and larger amounts of native EAAT2 protein. Proteomics data indicate a partial reversal of HD-induced changes in the EAAT2 interactor spectrum. Thus, astrocytic glutamate transport remains a target for therapeutic intervention.

## INTRODUCTION

HD is an autosomal dominant neurodegenerative disease. A trinucleotide (CAG) expansion of exon 1 in the mutant huntingtin gene *(mHTT)* causes abnormally long polyglutamine stretches in the already large huntingtin protein. *mHTT* is expressed not only in neurons but also in astrocytes (Faideau et al., 2010) where the loss of normal huntingtin function and additional effects of misfolded polyQ *mHTT* fragments can produce a large variety of alterations (Mrzljak and Munoz-Sanjuan, 2015;Zeitler et al., 2019;Tabrizi et al., 2019). HD patients suffer from progressive motor, cognitive and emotional disturbances (Tabrizi et al., 2013). A prominent motor symptom of advanced HD in humans and most rodent models of the disease is hypo-/bradykinesia (Berardelli et al., 1999;Hart et al., 2013;Horton et al., 2019), i.e. a reduced incidence and speed of self-induced movements. The appearance of jerky uncontrolled spontaneous movements (chorea) is the predominant feature at earlier stages of HD (Rosenblatt et al., 2003;Tabrizi et al., 2013). Thorough quantification of the hypo-*vs.* hyperkinetic aspects of motor performance is important for the pharmacological management of the disease (Hart et al., 2013). In general, phenotype progression is less faithfully predicted by the scores of chorea as compared to those of hypo-/bradykinesia (Rosenblatt et al., 2003;Tabrizi et al., 2013).

According to the information available from extracellular recordings or imaging of neuronal activity in intact HD mice, motor symptoms may evolve as a consequence of disinhibition and abnormal synchronicity in the corticostriatal pathway (Miller et al., 2012;Burgold et al., 2019). However, most studies point to functional uncoupling rather than enhanced glutamatergic input to the striatum (Plotkin and Surmeier, 2015;Reiner and Deng, 2018;Veldman and Yang, 2018). On the presynaptic side, there is a tendency for down-regulation of vGlut1 immunofluorescence (Rothe et al., 2015), decrease in vGluT1+ terminal numbers (Deng et al., 2013) and impairment of synaptic glutamate release at individual corticostriatal terminals (Dvorzhak et al., 2019). On the postsynaptic side, corticostriatal synaptic transmission might be affected by insufficient supply with brain-derived neurotrophic factor (Plotkin et al., 2014) or reduced signal transfer from distal dendrites (Carrillo-Reid et al., 2019). The duration of the NMDAR component of corticostriatal EPSCs was found to be prolonged (Dvorzhak et al., 2019), which might be due to a reduced glutamate uptake capacity of striatal astrocytes (Tong et al., 2014;Dvorzhak et al., 2016). Indeed, targeted expression of *mHTT* in astrocytes consistently impeded their glutamate uptake function (Shin et al., 2005;Bradford et al., 2009;Faideau et al., 2010;Meunier et al., 2016) and induced or exacerbated the motor symptoms of HD (Bradford et al., 2009;Meunier et al., 2016).

In the adult rodent striatum most of the glutamate transport is carried out by the excitatory amino acid transporter type 2 (EAAT2, gene name *Slc1a2)* (Danbolt, 2001;Beart and O’Shea, 2007;Vandenberg and Ryan, 2013). The EAAT2 protein is clustered at the sites of synaptic glutamate release (Danbolt, 2001) which ensures the rapid return of the extracellular glutamate concentration to very low resting levels (Bergles and Jahr, 1998). There is general agreement that *SLC1a2* expression is weaker in mouse models of HD. Moreover, spontaneous and/or pharmacologically stimulated uptake of exogenous [3H]-labeled L-glutamate or D-aspartate was reduced by HD, as shown in lysed synaptosomal preparations (Lievens et al., 2001;Bradford et al., 2009;Faideau et al., 2010;Huang et al., 2010;Parsons et al., 2016) or striatal perfusates (Behrens et al., 2002;Estrada-Sanchez et al., 2019). Nevertheless, a number of studies have challenged the idea that reduced expression of *Slc1a2* in HD astrocytes would actually translate into a state of deficient synaptic glutamate clearance (Parsons et al., 2016;Parievsky et al., 2017) or contribute to the progression of HD (Petr et al., 2013). Obviously, more information was needed to clarify the conditions resulting in the proposed mismatch between glutamate release and clearance.

In principle, a relative deficiency of glutamate uptake could result from the following distinct mechanisms: i) altered transporter-substrate interactions, ii) insufficient *Slc1a2* transcription, iii) impaired membrane targeting of EAAT2, and/or iv) exaggerated intracellular degradation of EAAT2. Heterologous expression studies with recombinant chimeric EAAT2 variants delineated the significance of functional compartments within the protein. It turned out that a region close to the distal end of the last transmembrane domain is relevant for the interaction with the transport substrate (Leinenweber et al., 2011), and affects the balance between cytoplasmic retention and membrane insertion (Kalandadze et al., 2002;Underhill et al., 2015). Due to detailed studies from the Zafra lab, it is also known that ubiquitination of 4 C-terminal lysine sites (Gonzalez-Gonzalez et al., 2008) substantially contributes to the internalization of the transporter from the plasma membrane. Finally, it was reported that sumoylated toxic C-terminal fragments may inhibit the *Slc1a2* transcription in the astrocyte nucleus

(Gibb et al., 2007;Foran et al., 2011;Rosenblum et al., 2017). All these studies imply that EAAT2 binding partners at the C-terminal end participate in the regulation of glutamate uptake, but detailed information on the spectrum of EAAT2 interactors in health and disease is not yet available.

The present study addresses the possibility that glutamate uptake can be enhanced by the expression of genetically engineered variants of *Slc1a2.* If so, this would validate the proposed links between EAAT2 structure, glutamate uptake activity, synaptic glutamate clearance and exploratory motor behavior.

## RESULTS

### Experimental approach

Fig. 1a, b illustrates the principal elements of the current series of experiments on modified *Slc1a2* expression. All data was obtained from aged (14-18 mo) wild-type (WT) and heterozygous (HET) Q175 mice, a well established mouse model of HD (Menalled et al., 2012). Adeno-associated viral vectors were used for the transfer of recombinant DNA into the dorsal striatum, the region most affected by HD-related neurodegeneration (Vonsattel and DiFiglia, 1998). The respective plasmids were derived from human EAAT2, with the understanding that there is a close similarity between the human and mouse *Slc1a2* homologues (Kirschner et al., 1994). For detection of the transgene, mRuby3 or, in some experiments, mYFP was fused to the N-terminus of the *Slc1a2* sequence. The CONTROL groups (WT-CTRL, HET-CTRL) received a vector solely encoding the fluorescent tag. The TEST groups (only HET mice) received a vector encoding one of 3 EAAT2 variants. The TEST group of HET-EAAT2-S506X expressed a strongly modified EAAT2 variant with removal of the last 68 amino acids, i.e. almost complete truncation of the C-terminal domain. Mice belonging to the TEST group of HET-EAAT2-4KR expressed EAAT2 with four point mutations at the C-terminal domain. The mice of the third TEST group expressed the full-length EAAT2. Thus, the 3 TEST groups represented three degrees of C-terminal modification, from almost complete truncation to full-length EAAT2. HET-CTRL is the principal reference group for comparison with WT-CTRL (to characterize the HD-phenotype) and HET-TEST (to determine the effect of treatment).

**Fig. 1.**
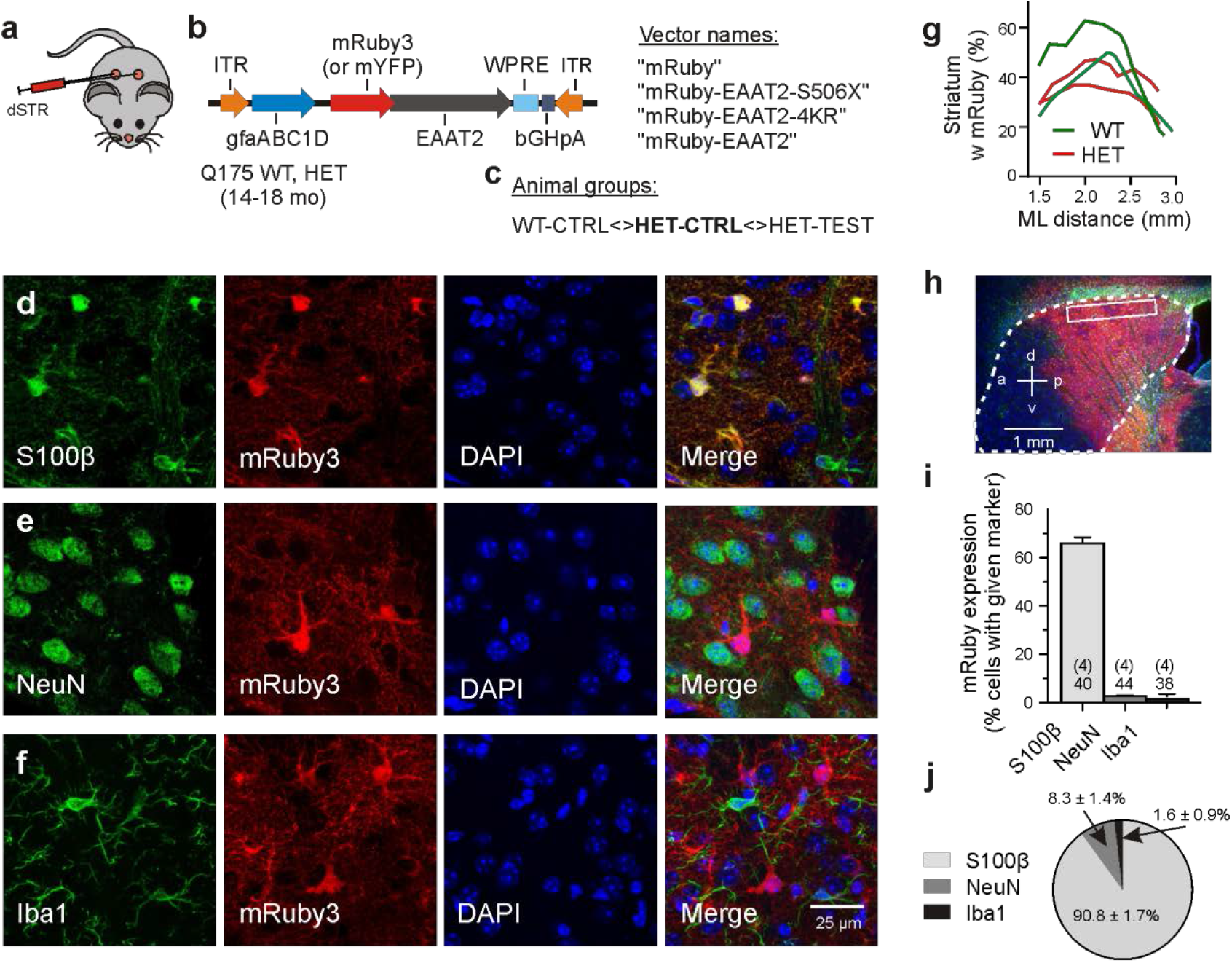
Experimental approach. **a, b** Mouse model and types of injected vectors. **c** General test scheme. **d-f** Immunostained sectionsfor evaluation of viral expression patterns. Triple labeling with representative astrocytic (d), neuronal (e) or microglial (J) markers, together with an antibody against mRuby3 and nuclear counterstaining with DAPI. **g** The mRuby+ area at the indicated mediolateral distance. **h** Transduced area visualized with mRuby3 on the background of DAPI (blue) and GFAP immunofluorescence (green). The putative margin of the striatum is marked with a dotted line. **i** Incidence of transduced cells within the S100β-, NeuN- or Ibal-expressing cell populations. Numbers on columns: Evaluated sections and animals (in brackets). **j** Fraction of the astrocytic, neuronal or microglial phenotype among the mRuby3 + population in the dorsal striatum. Abbreviations: dSTR - dorsal striatum. M1-CTX - primary motor cortex (M1).

The expression of the transgene was limited by the chosen experimental conditions, including i) the amount of injected particles (a single injection of 1.0*10^9^gc/striatum), ii) the expression time (3-5 weeks, if not mentioned otherwise), iii) the type of the viral serotype (PHP.eB) and iv) the promoter sequence (gfaABC1D). To examine the abundance and specificity of transduction we analyzed immunostained parasagittal sections from fixed brains (Fig. 1d-f). As functional imaging experiments were limited to cells in the dorsal striatum, counts were performed at a mediolateral distance of 2 - 2.5 mm within the boxed area. Fig. 1g, h illustrates that the area of evaluation was always well within the extensively labeled zone. The latter covered about 40% of the striatal volume. An “effective transduction rate” (ETR) was determined in 4 mice (2 WT and 2 HET), each contributing about 10 viewfields. Co-localization of mRuby3 with a suitable cell-specific marker rendered the following ETR values (Fig. 1i): S100β+ cells (astrocytes) - 65.9 ± 2.47%, NeuN+ cells (neurons) - 2.76 ± 0.44%, Iba1+ cells (microglia) - 1.64 ± 0.85%. There also was some labeling in the overlaying cortex and a small degree of unintended transgene expression in neurons and microglial cells. Among the population of mRuby+ cells in the dorsal striatum (boxed area in Fig. 1h), 90.8% were astrocytes, 8.3% - neurons and 1.6% - microglial cells (Fig. 1j).

The cell counts indicate that the viral transduction was local (dorsal striatum), specific (astrocytes) and effective (two thirds of the astrocytes in the labeled area). This expression model was therefore considered suitable to characterize the HD phenotype, on one side, and the response to the exogenous EAAT2 variants, on the other side.

### Motor performance of control and treated HD mice

HET mice exhibit the symptoms of hypokinesia at an age of 10-12 months meaning that at the time of injection and testing neurons and astrocytes in the striatum and elsewhere have undergone changes for several months. The aim of the following experiments was to quantify the HD-induced changes in locomotion and to find out to what extent these changes could still be reversed. To gain a set of easy-to-quantify indicators of hypokinesia the mice were submitted to two video-recorded tests: i) the step-over latency test, SOLT (Dvorzhak et al., 2019) and ii) the open field test, OFT (Menalled et al., 2012;Rothe et al., 2015). The tests were performed immediately before the viral vector injection and 3-5 weeks later, before animal sacrifice. This behavioural examination provided us with a set of 6 criteria to classify an animal or animal group as being hypokinetic or recovered from hypokinesia.

Specifically, the step-over latency of SOLT was regarded as a direct measure of the time needed to initiate exploration in the open field. Its usefulness was supported by a larger number of observations from non-injected HD mice (Supplemental Fig. 1). Step-over latencies longer than 300 ms were almost exclusively found in HET which could contribute to the detection of recovery effects. The other indicators are based on a more complex analysis of the movement trajectories in the open field (Fig. 2a, b, see Supplemental text for definitions of the analysed parameters). In the present experiments with the injected WT-CTRL, HET-CTRL and HET-EAAT2-S506X TEST mice, 5 out of 6 movement parameters extracted from OFT were sensitive to the C-terminal-modified EAAT2. The most commonly used OFT indicator “total distance travelled in 5 min” was found to be decreased in the HET-CTRL group and increased in HET-S506X if compared to HET-CTRL (Fig. 2a, c). The incidence of starts from rest is possibly the most reliable indicator of depressed locomotion (Fig. 2a, d). It is inversely correlated with the step-over latency (Fig. 2f). After treatment with mYFP-EAAT2-S506X both parameters exhibited changes towards WT levels (Fig. 2d, f). Also noteworthy is the treatment-related reduction of the maximal to mean radius of the resting area, i.e. the extension of the resting area until a new movement starts (Fig. 2b, g). One could further expect that HD or treatment can alter the velocity of body displacement, but it is not known whether this would equally apply to rest and running. Previous open field experiments with simultaneous recording of motor activity and local field potentials (LFPs) suggested that the HD-related synchronisation of striatal activity only occurs during the phases of rest thereby underlining the distinct character of the resting condition (Rothe et al., 2015). Now it turned out, that movement velocity is affected both during the resting and running phases. Moreover, the direction of the changes induced by HD and treatment were the same (Fig. 2h, i). Other aspects of open field behavior were unchanged by viral treatment, such as the relative time spent in the open field center, or the open field resting time (Fig. 2j, k) which suggests that these indicators might be unrelated to the site of injection, the dorsal striatum, or not be representative of exploratory motor activity.

**Fig. 2.**
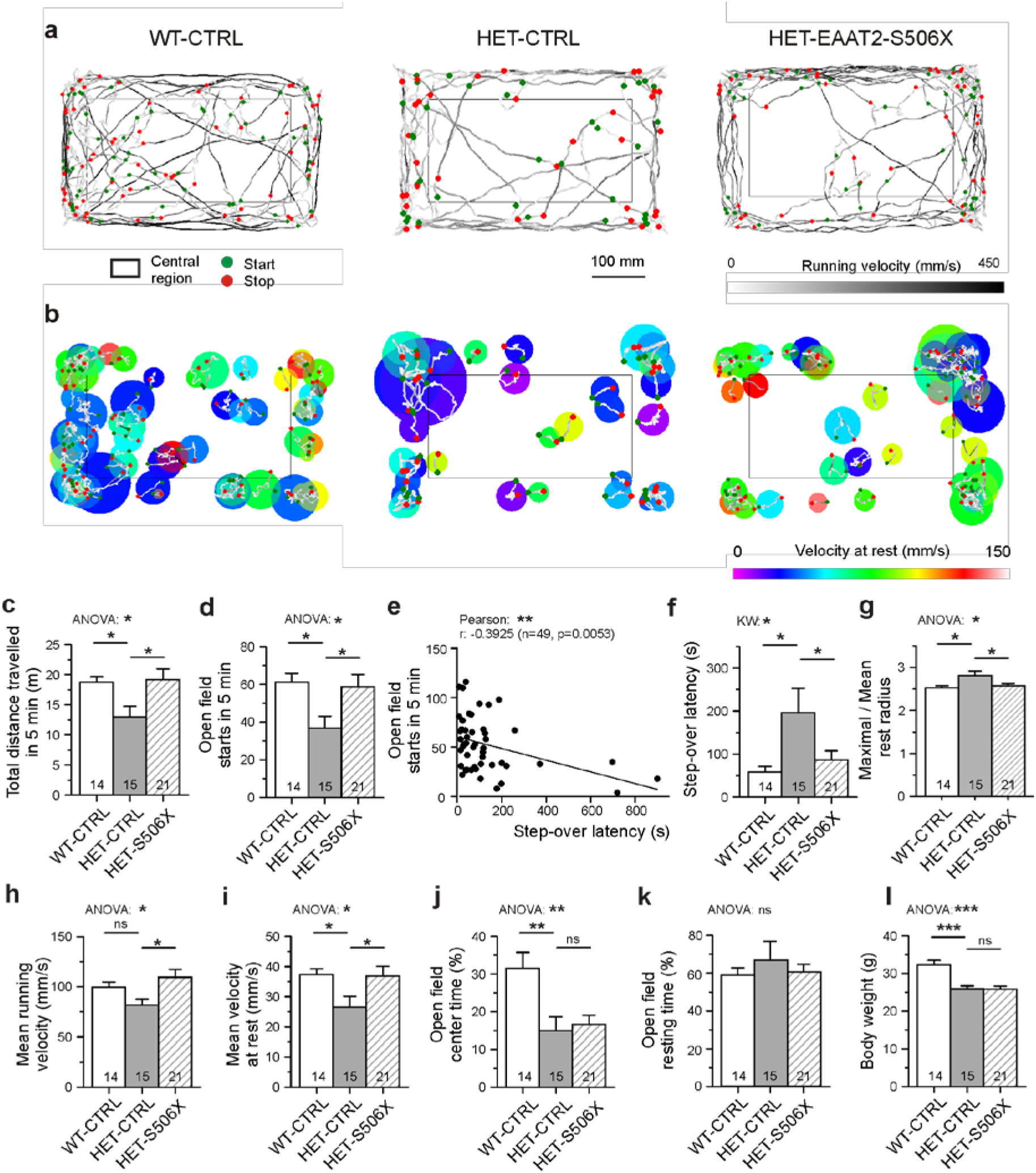
Effects of bilateral EAAT2-S5O6X expression on the motor performance of QI 75 mice. **a** Recordings of open field trajectories showing the mean running velocity between starts and stops (green or red circles, respectively). Gray level coding of instantaneous running velocity. **b** Motor activity at rest. Color coding of movement velocity at rest. **c, d, f-k** Quantification of results from open field testing. **e, f** Inverse relationship between the number of starts from rest and step-over latency, and sensitivity of the latter to HD and treatment. **l** matched age-, CAG- and body-weigfit composition of the test groups (only body weight illustrated).

Among the less effective attempts to return the functional indicators of HET to WT performance were experiments with mice expressing a full-length EAAT2 variant. In this case the enhancement of locomotion was small, if at all present (compare the corresponding graphs in Fig. 2c-i and Supplemental Fig. 2a-e). The statistical analysis showed that EAAT2-S506X-treated mice outperformed not only with respect to HET-CTRL but also with respect to HET-EAAT2. This was significant (ANOVA p<0.05) for the number of open field starts stops in 5 min and the mean running velocity (mm/s). The respective mean, SE and N were for HET-CTRL: 36.80 ± 6.23 and 81.57 ± 6.04 (N=15), HET-EAAT2-S506X: 58.67 ± 6.66 and 109.45 ± 8.06 (N=21), HET-EAAT2: 41.40 ± 7.38 and 94.45 ± 8.57 (N=10).

Together, these experiments suggest i) that the state of striatal astrocytes is relevant for the incidence and speed of exploratory movements, ii) truncation of the EAAT2 C-terminal might be an effective means to reduce locomotor deficits in HD mice and iii) induced over-expression of full-length *Slc1a2* is not sufficient to alleviate the symptoms of hypokinesia.

### Astrocytic glutamate uptake

To gain further information on the functional consequences of EAAT2 modifications in HD, imaging experiments were performed in striatal slices loaded with the Na^+^ indicator SBFI-AM. Glutamate transport was elicited with L-aspartate. It is known that L-glutamate, L-aspartate D-aspartate are taken up with similar micromolar affinity (Arkhipova et al., 2019). In the present experiments L-aspartate was chosen for its physiological role as excitatory neurotransmitter (Morland et al., 2013) and its negligible effects at G-protein-coupled glutamate receptors. The major part of the L-aspartate-induced response was blocked by the high-affinity glutamate transport inhibitor TFB-TBOA (Fig. 3a), and this TFB-TBOA-sensitive component of the Na^+^ elevation was quantified in WT-CTRL, HET-CTRL and two HET-TEST groups, HET-EAAT2-S506S and HET-EAAT2-4KR. The recordings of averaged responses (Fig. 3b) visualize the differences between WT and HET and the similarity of responses obtained in the WT-CTRL, EAAT2-S506X and EAAT2-4KR groups. Multilevel (nested) ANOVA confirmed a significant rescue of the L-aspartate-induced sodium response in astrocytes transduced with EAAT2-S506X and EAAT2-4KR (Fig. 3c and Supplemental Tab. 1). The present SBFI-loading protocol was adjusted to preferentially label astrocytes (Dvorzhak et al., 2016). This offered an opportunity to determine, in the same viewfield, the L-aspartate-induced glutamate uptake activity of transduced versus non-transduced astrocytes (Fig. 3d, e). In the case of HET-EAAT2-S506X, there was a significant difference between the transduced and not-transduced astrocytes (Fig. 3f).

**Fig. 3.**
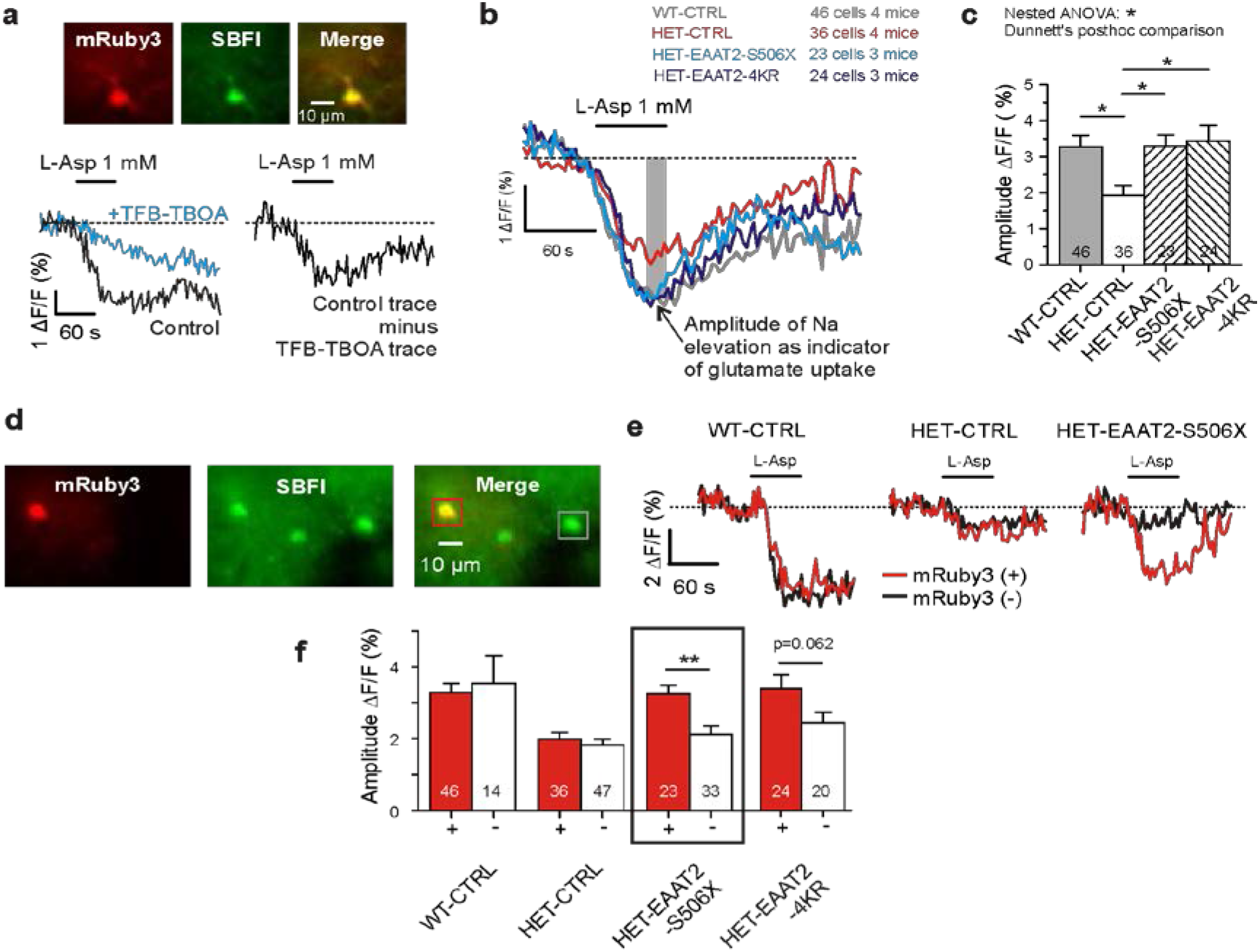
Rescue of glutamate uptake activity of striatal astrocytes after bilateral expression of EAAT2-S5O6X or EAA T2-4KR. **a** Measurement of glutamate uptake by sodium imaging with SRFĨ. Traces recorded in the absence and presence of TFB-TBOA (2 μM). The amplitude of this differential response was expressed as Δ*F/F* during the last 15 sec of L-Asp application. F is the mean fluorescence at rest, before drug application. **b** Averaged traces of SBFI fluorescence. Results from mRuby3·positive astrocytes only. **c** Quantification of the L-aspartale-induced sodium elevation. Two-level (“nested”) statistics (animal level, cell level and posthoc comparison between groups). **d** The applied SBFI loading protocol was selective for astrocytes. Vìewfìeld with 3 astrocytes, where one is transduced and two are not. **e** Traces from mRuby+ as opposed to mRuhy-in three test groups. Averaged traces from one. animal per group. **f** Comparison of results obtained in the different groups from mRuby3 + *vs.* mRuhy3-astrocytes.

SBFI imaging was also performed in mice injected with the vector encoding full-length EAAT2 (Supplemental Fig. 2f, g), but significant rescue of glutamate uptake was not observed. Statistical analysis of the L-aspartate-induced Na transients in HET-CTRL, HET-EAAT2-S506X and HET-EAAT2 (Kruskal-Wallis test: P=0.0004) showed that EAAT2-S506X-treated mice outperformed not only with respect to HET-CTRL (Dunn’s test: P = 0.0004) but also with respect to HET-EAAT2 (Dunn’s test: P = 0.0052). The respective mean, SE and N of ΔF/F (%) were for HET-CTRL: 2.01± 0.18 (N=36), HET-EAAT2-S506X: 3.28 ± 0.23(N = 23), HET-EAAT2: 2.31 ± 0.23 (N = 34).

These results show that the expression of the truncated but not the full-length EAAT2 variant effectively counteracted the HD-induced depression of glutamate uptake.

### Possible effects on neuronal excitability and results of systemic vector application

A series of supplemental experiments addressed the influence of HD and EAAT2-S506X treatment on neuronal excitability (Supplemental Fig. 3) and the efficacy of treatment with systemically applied vectors (Supplemental Fig. 4). In both cases the effects of S506X remained below the significance level required with nested ANOVA. These experiments may however provide useful information on the HD-related pathologies in striatal astrocytes of aged mice.

### Glutamate clearance at corticostriatal synapses

The significance of astrocytic glutamate uptake deficits for glutamatergic synaptic transmission in the striatum is still a matter of debate. A previous publication from our lab (Dvorzhak et al., 2019) has addressed this controversy by using CaMKII-driven expression of the genetically encoded “ultrafast” glutamate indicator iGlu*u* (Helassa et al., 2018) to examine the performance of single corticostriatal synapses in WT and Q175 HET. It was found that in HET aged 15 to 19 months at least 40% of corticostriatal synapses exhibited a deficit in the glutamate clearance. A rescue of the synaptic glutamate clearance function in symptomatic HD mice after a molecular intervention targeting the astrocytic glutamate uptake would be a strong argument in support of the clearance-deficit hypothesis of HD. We therefore examined the effect of the truncated EAAT2-S506X transgene expression on the properties of single corticostriatal synapses in a group of HD mice with manifest symptoms of hypokinesia.

Large (>0.63 μm) fluorescent terminals were assumed to belong to the group of PT-type afferents formed by the layer V pyramidal neurons of the motor cortex (Reiner and Deng, 2018;Dvorzhak et al., 2019). The specimen recordings selected for Fig. 4a illustrate the differential spread and duration of the glutamate elevation in the perisynaptic space. Only terminals located on the territory of transduced astrocytes were included in the test samples. The arrowhead of Fig. 4b points to a bright varicosity on the background of the red fluorescent area generated by the dendritic field of one mRuby-expressing astrocyte. These spatial interrelations are relevant for a better understanding of synaptic modulation by astrocytic transporters or gliotransmitters.

**Fig. 4.**
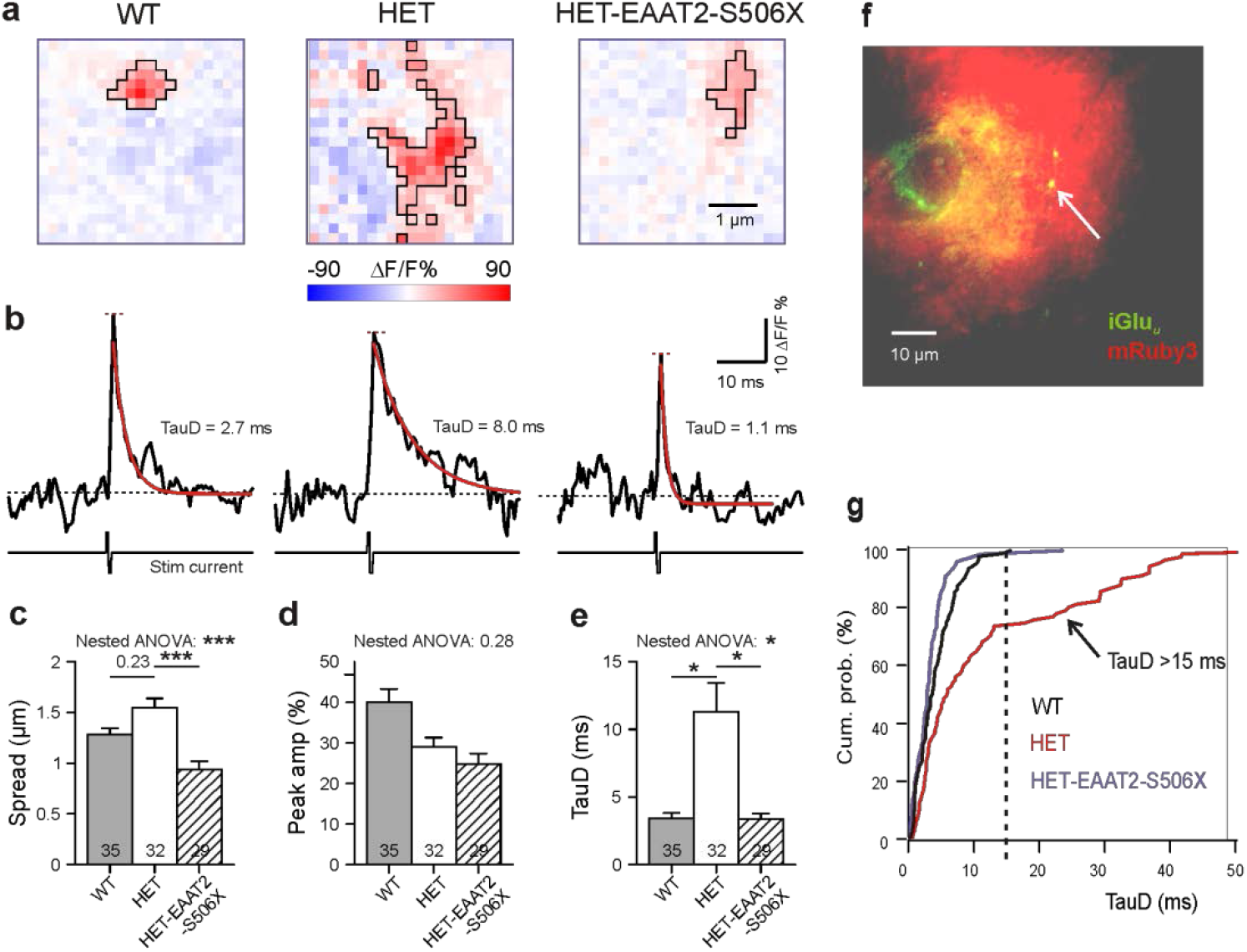
Rescue of single synapse glutamate clearance in the striatum after expression of EAAT2-S5O6X. **a** Specimens from single PT-type coríicostriatal synaptic terminals in siices from WT HET and treatedHET. Bilateral intracranial injections of HET with EAAT2-S5O6X. The selected images were acquired at the response peak to illustrate the extension of iGlu_L·_ elevation (boxed area). Pink pixels within the black boundaries are pixels where stimulation of the bouton elicited a fluorescence increase (ΔF) to values larger than the resting level prior to stimulation (F). **b** Corresponding iGlu_u_ values before and after electrical stimulation of the given bouton in the presence ofTTX. The traces represent the mean value calculatedfrom all suprathreshold pixels in the ROT Red curves: Monoexponential functions fitted to the iGlu_u_ traces. In black - respective time constants of decay (TauD). **c-e** Quantification of results. **f** Only synapses in the immediate vicinity of transduced astrocytes were included, as confirmed by the location of the tested iGlu_u_-expressing varicosity on the territory of mRuby3-positive astrocytes (arrowhead). **g** Cumulative probability plot to illustrate the absence of TauD values >15 ms in WT and in EAAT2-S5O6X-treated HET. TauD values >15 ms identify pathological synapses in HET.

It can be seen that the parameter “Spread” (see Methods for definition) was sensitive to treatment with EAAT2-S506X (Fig. 4a, d). Fig. 4c shows the time-dependent changes of the iGlu*u* intensity values before and after a short biphasic electrical stimulus applied through a glass pipette to the iGlu*u*-expressing presynaptic terminal. Such traces could be used to extract two important indicators: “Peak amplitude” (short red horizontal line) and “Time constant of glutamate concentration decay” (TauD, see values next to the red monoexponential fitting curve). Similar traces were also constructed from single pixels. The transients derived from the pixel with the highest fluorescence increase reflect the release and clearance of glutamate next to the presynaptic active zone, i. e. the site of vesicle exocytosis. The respective results are given in the Supplemental Tab. 1.

The data presented in Fig. 4d-f is based on the mean suprathreshold pixel intensity values. It can be seen that Spread and TauD, but not Peak amplitude, were affected by the expression of EAAT2-S506X. It is, thus, very likely that Spread and TauD reflect the astrocytic glutamate uptake while the Peak amplitude indicates the amount of released glutamate (Dvorzhak et al., 2019). TauD values larger than 15 ms have never been observed in WT while more than one third of the HET synapses tested at an age of ~16 months showed TauD values >15 ms. Now we found that in HET treated with EAAT2-S506X synapses with prolonged decay were entirely absent (Fig. 4g).

These results demonstrate i) a contribution of the astrocytic glutamate uptake deficit to the HD-related pathology in the corticostriatal pathway and ii) a return of synaptic glutamate clearance towards wild-type levels after expression of the C-terminal-truncated EAAT2 variant.

### Native intrinsic EAAT2 in immunostained corticostriatal synapses

Next it was asked whether effects of EAAT2-S506X (or EAAT2-4KR) could also be discerned in EAAT2-immunostained sections. It should be noted that the selected antibody (Ab) only reacts against EAAT2 with intact C-terminal. This provided us with an opportunity to examine the effect of transgene expression on the native perisynaptic EAAT2. Fig. 5 shows the results from four simultaneously processed test groups, each consisting of two CAG-, age- and body weight-matched animals: WT-CTRL, HET-CTRL, HET-EAAT2-S506X and HET-EAAT2-4KR. The images obtained from the immunostained sections (Fig. 4a) illustrate, first of all, a noticeable shift towards lower EAAT2 intensity levels in HET, thereby confirming previous studies from non-injected HD mice (Estrada-Sanchez et al., 2009;Tong et al., 2014). Fig. 5b shows the respective single synapse immunofluorescence (IF) in ROIs containing only one vGluT1-positive spot, i.e. one corticostriatal presynaptic terminal. The EAAT2-positive pixels co-localized with mRuby3, i.e. they belongrd to an astrocyte process. The graph of Fig. 5c and the Supplemental Tab. 1 show the results of statistical evaluation of the EAAT2-immunofluorescent clusters within the analyzed ROIs. Both the HET-EAAT2-S506X and the HET-EAAT2-4KR groups differed from the HET-CTRL group, as their synaptic EAAT2 protein level increased to nearly WT level.

**Fig. 5.**
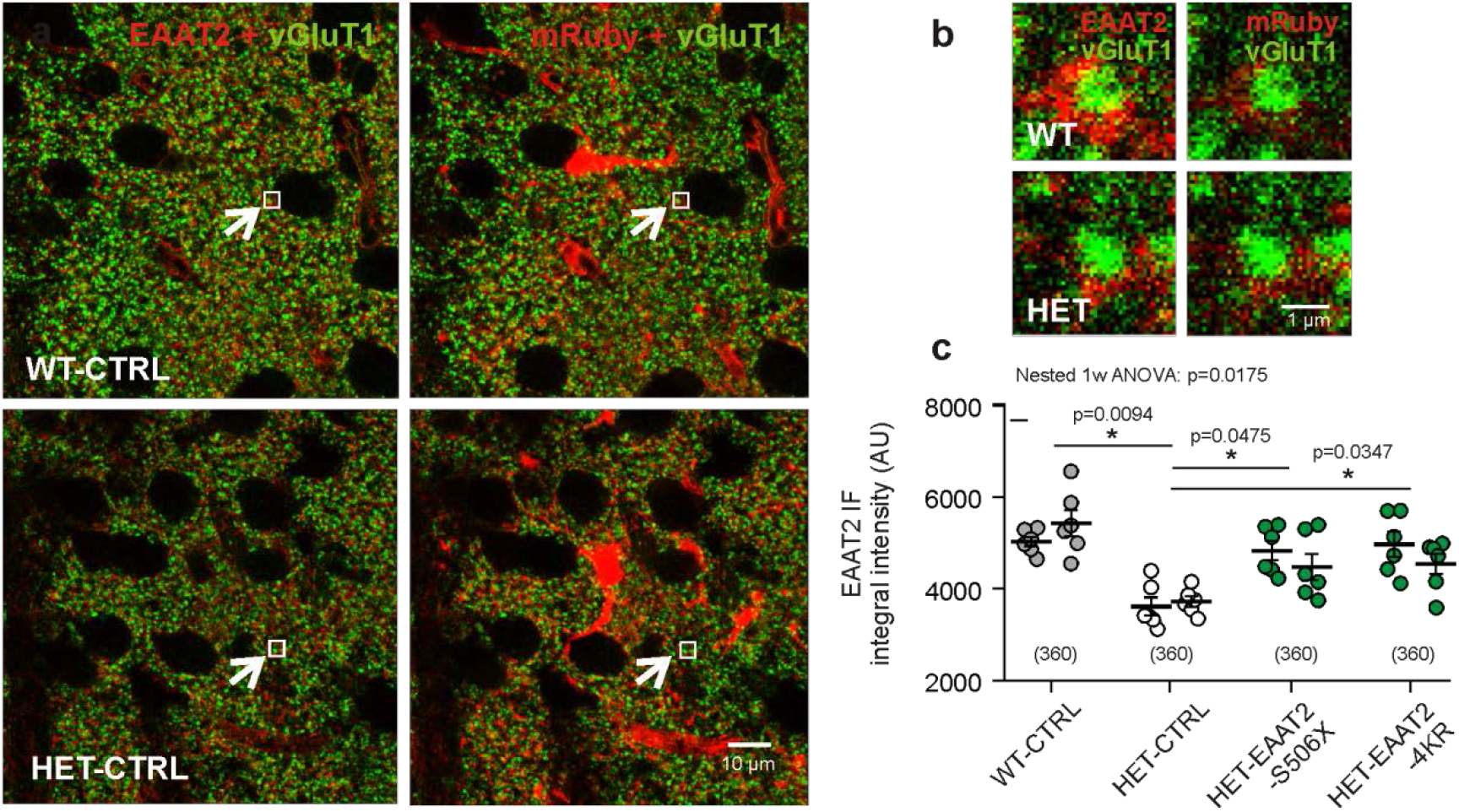
Evaluation of synaptic EAAT2 immunofluorescence in fixed sections. **a** Confocal images from WT-CTRL and HET-CTRL to illustrate the HD-related decrease in the overallEAAT2 IF. In the illustrated samples the mean intensity values were 3078 ± 38.4 (WT-CTRL) and 2673 ± 31.7 (HET-CTRL). **b** Regions of interest (ROIs) comprisingjust one synapse. The ROIs correspond to the small boxed areas in **a**. The EAAT2~immunopositive area corresponds to an astrocyte PAP, as it is labeled with mRuby3. **c** Quantification of integral EAAT2 IF from ROIs comprising only one vGluTl-immunopositive terminal. Numbers in brackets: evaluated synapses per group. Each data point is the mean valuefrom 30 synapses on 1 mRuby3-labeled astrocyte. 12 astrocytes per groupfrom 2 mice/group. Statistics: Nested data analysis.

The data from immunostained brain sections suggest an effect of the C-terminal-modified EAAT2 on the level of the native murine EAAT2 at corticostriatal synapses.

### Western blot (WB) analysis of native EAAT2

The following experiments were designed to further examine the possibility that expression of a C-terminal-modified EAAT2 transgene in the HET striatum can increase the amount of native EAAT2. About 15 months old Q175 WT and HET were sacrificed 3-4 weeks after viral injection to prepare striatal lysates for WB analysis. The lysates contained both the “foreign” YFP-tagged EAAT2 transgenes and the “intrinsic” native EAAT2 protein without tag (Fig. 6a). The compared animal groups are listed in Fig. 6b. An overview of all possible WB bands is given in Fig. 6c. The native EAAT2 monomers and dimers are represented in the 70 kDa and 140 kDa bands, respectively. Due to the C-terminal binding characteristics of the EAAT2 Ab, immunoreactivity (IR) is only observed at the native EAAT2 and YFP-EAAT2 bands but not the C-terminal truncated YFP-EAAT2-S506X band (compare boxed area in Fig. 6d and e). Evaluation of the striatal lysates from HET-EAAT2-CTRL and HET-EAAT2-S506X showed that the amount of native EAAT2 was indeed higher in the HET-EAAT2-S506X group (Fig. 6f) which lends further support to the suggestion that the transgene and native EAAT2 proteins interact.

**Fig. 6.**
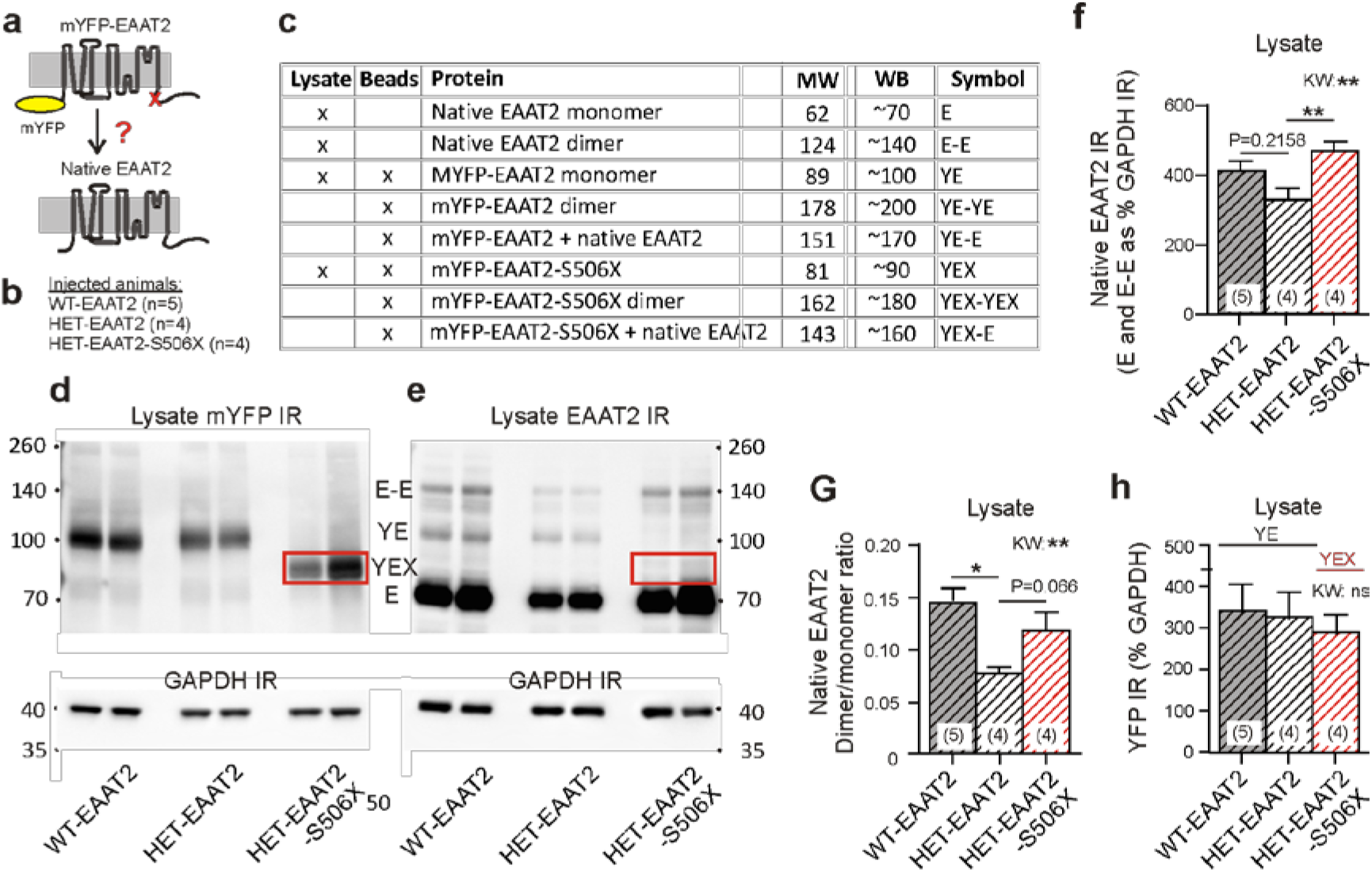
Western blot (WB) analysis to examine the changes in native murine EAAT2. **a** Two principal forms of EAAT2 to be detected in the lysates: Native murine EAAT2 and recombinant YFP-EAAT2 expressed due to virus injection. **b** Test groups according to injected mice. **c** EAAT2 bands to he delected in the WB. Info for the immunoprecipitate (beads) to be used for Fig. 7h. **d, e** WB samples prepared for the quantification of YFP and EAAT21R in striatal lysates. At leastfour animals per group (shown are two). The boxed band YEX is not detected in **e** since the preferred antibody against EAAT2 binds to a region located on the del eted C-terminal. **f** Mean intensity of native EAAT21R (mono- and dimers). Note significantly higher amount of native EAAT2 in HETs expressing the truncated as opposed tofull-length EAAT2 variant. **g** Quantification of the dimer to monomer ratio E-EE. **h** Similar amounts of mYFP 1R (YE or YEX bands) expressed in the 3 animal groups. Nonparametric Kruskal-Wallis test (KW) and Dunn’s posthoc comparison applied in all 3 graphs.

It is conceivable that the smaller amounts of synaptic EAAT2 in HD were associated with a deficit in multimer formation. Under the given experimental conditions, i.e. Western blotting, only a small percentage of EAAT2 (less than 15%) retained or re-established a dimeric form. Nevertheless, it could be shown that the dimer/monomer ratio of native EAAT2 was lower than in the HET-EAAT2 as compared to the WT-EAAT2 group. But the treatment effect remained below significance level. Of note, the amount of the mYFP-tagged transgene was not different between the test groups (Fig. 6h).

Together with the data in Fig. 5, these findings provide a first hint on the mechanism underlying the behavioral and cellular rescue effects in HD mice treated with the truncated EAAT2 transgene. The results are consistent with the idea that expression of the S506X variant can increase the availability of native EAAT2.

### Proteomics analysis of the EAAT2 interactor spectrum in the striatal immunoprecipitate

Further understanding of the signaling pathways underlying the regulation of astrocytic glutamate transport would require a better knowledge of the involved EAAT2 binding partners. Such information could be obtained from massspectrometry-based analysis of the EAAT2 interaction proteome after a “pull-down” of YFP-tagged EAAT2. Fig. 7a illustrates the selected method of immunoprecipitation from striatal lysates where nanobody-coated magnetic beads served as “bait” for the YFP fused to the EAAT2 protein. The animal groups were the same as in Fig. 6: WT-EAAT2, HET-EAAT2 and HET-EAAT2-S506X. The quality of the immunoprecipitate was verified by WB analysis of YFP- and EAAT2-IR (Fig. 7b, c).The total amount of YFP pulled down by the beads was very similar in the 3 tested groups (Fig. 7d), as should be the case if the amounts of “bait” were equal in the 3 groups, and the amount of “prey” (YFP-EAAT2) would have sufficed to saturate the binding sites.

**Fig. 7.**
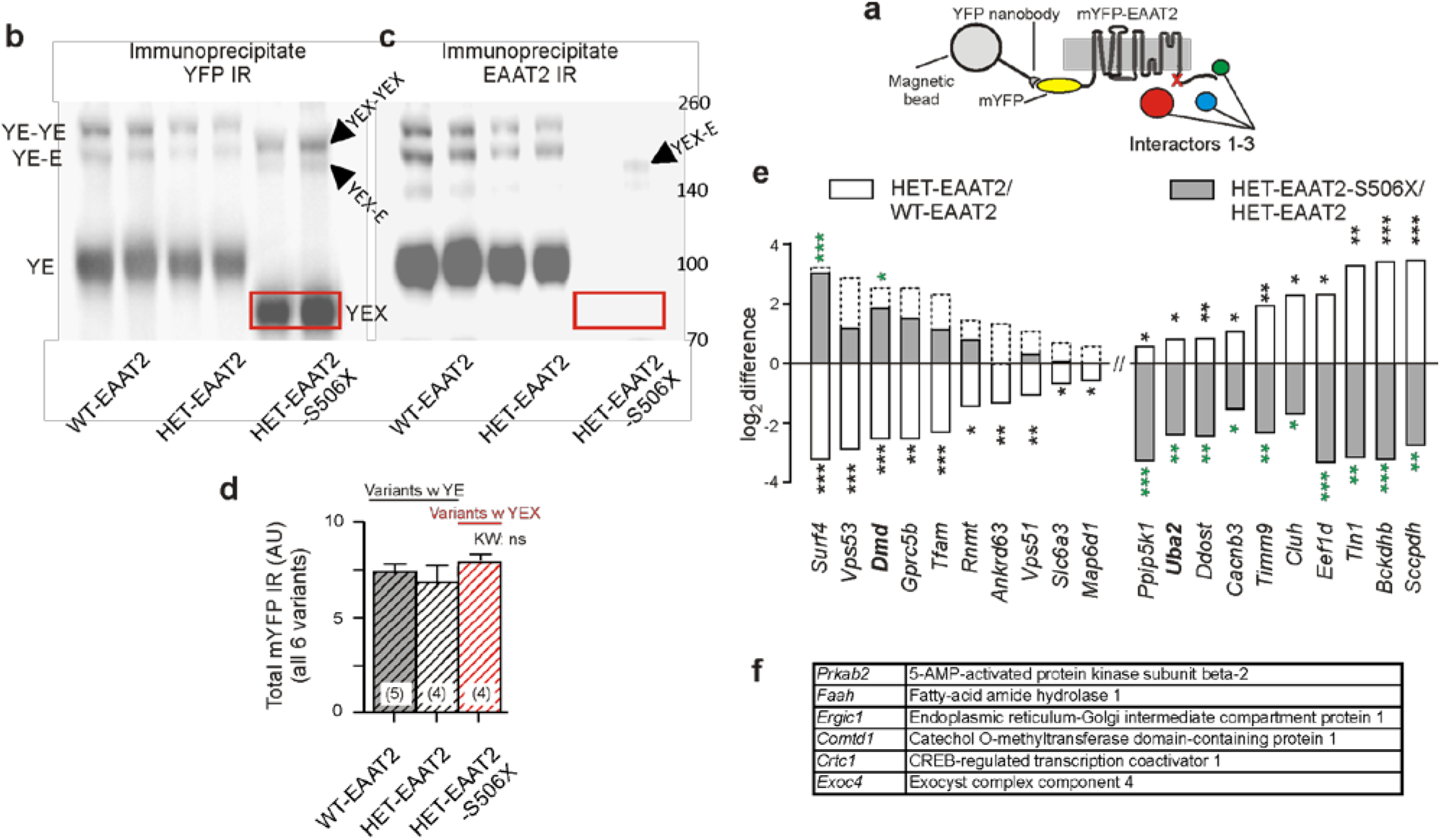
HD- and treatment-related changes in the abundance of mYFP-EAAT2 binders. **a** Scheme of experiment. **b, c** WB samples preparedfrom the YFP-immunoprecipitate. For the identification of bands see the table in Fig. 6b. Note more prominent dimer bands in comparison with the WB samples prepared from lysates. **d** Similar amounts of m YFP IR (YE or YEN bands) pulled down in the 3 animal groups. **e** Quantification of HD- and treatment-related changes in the abundance of the mYFP-EAAT2 binders. Data from LFQ quantification. The same binders were found with iBAQ quantification. Only proteins with >3 peptides detection and abundance of log2 intensities > = 23 in all 4/5 animals ofa group were included. The signalfrom each animal was normalized to the mYFP median intensity value in a histogram constructed from all samples. The inter-group intensities were compared by a two-tailed t-test. Empty bars: log2 difference (or FC ratio) of binder amounts in the HET-EAAT2 vs WT-EAAT2 groups. Bottom line: List of 20 interactors with significant HD-related difference (black asterisks on empty bars). An HD-related down-regulation is shown as downward bars, and an up-regulation - as upward bars. The same interactors were examined for changes between the HET-EAAT2-S506X vs. HET-EAAT2 groups (filled bars). In this case a significant change is denoted by green asterisks. The dashed bars denote the level required to reach WT levels. **f** Identification of 6 significant (p<0.01) EAAT2 interactors present both in WT and HET but without effect of HD or C-terminal truncation, p<0.05 (*). p<0.01 (**) and p <0.001

The immunoprecipitate was then submitted to liquid chromatography tandem mass spectrometry (LC-MS) for further analysis of the EAAT2 C-terminal binder spectrum (Fig. 7a). The following experiments were designed to compare i) the amount of full-length EAAT2 interactors in HET *vs.* WT and ii) the amount of truncated *vs*. full-length EAAT2 interactors in HET. Each experiment was performed in at least 4 mice, with 2 repetitions per animal. Results not confirmed by a 2nd run were discarded. Fig. 7e lists 20 EAAT2 interactors arranged according to the size/direction of the HD-related differences. Significant differences in the abundance of a given interactor are denoted by asterisks. The empty bars/black asterisks reflect the differences between the WT-EAAT2 and HET-EAAT2 groups, i.e. the effects of HD as such. The filled bars/green asterisks show the respective log2 differences between the HET-EAAT2-S506X and the HET-EAAT2 groups. This data illuminates the impact of C-terminal truncation and may include rescue effects. Please note that the experiments cannot show what actually happens to the native murine EAAT2 protein devoid of mYFP, but rather provide a first glimpse at the normal and pathological EAAT2 binding partners that may influence the glutamate uptake function in striatal astrocytes of intact mice. It can, for instance, be seen that the interaction with dystrophin (gene name *Dmd)* is decreased in HD and partially restored in HET treated with EAAT2-S506X. Most interesting, HD is also associated with a substantial up-regulation of potential EAAT2 interactors, an example being UBA2, the ubiquitin-like 1-activating enzyme E1B, also known as SUMO-activating enzyme subunit 2 with the gene name *Uba2.* This interaction is strongly reduced by the truncation of the C-terminus in the HET-S506X group.

Other highly abundant or significant interactors of EAAT2 were not affected by HD (Fig. 7f) or not covered by the test. The present assay with a unique peptide number of >3 included 3.790 from a total of 13.000 proteins so far identified in the mouse brain (Sharma et al., 2015).

These proteomics results are consistent with the hypothesis that: i) the EAAT2 interactome is influenced by HD, ii) some of the HD-related changes are reversible and iii) expression of C-terminal-truncated EAAT2 may block abnormal interaction in favor of more physiological protein interaction patterns.

## DISCUSSION

The results obtained with intrastriatal injection of viral EAAT2 constructs support the hypothesis that HD-related deficits can be alleviated by the expression of a C-terminal-modified EAAT2 transgene. This conclusion is based on changes in self-induced locomotion, astrocytic glutamate uptake, synaptic glutamate clearance and synaptic EAAT2 protein levels. Together, the results underscore the importance of C-terminal-dependent EAAT2 binding partners in HD-related pathologies.

### Is region- and cell-type-restricted expression of artificial EAAT2 variants a useful approach to study disease-related pathology?

HD is a severe inherited neurological disorder with diverse clinical symptoms and variable onset. With a respective gene test it can be diagnosed much before the onset of motor or other symptoms. Great efforts are therefore being made to reduce the expression of *mHTT* inside and outside the central nervous system before the disease actually starts (Mrzljak and Munoz-Sanjuan, 2015;Tabrizi et al., 2019). *mHTT*-lowering therapies have reached an advanced stage of preclinical testing or even entered the phase of first clinical trials (Mullard, 2019). In animal models of HD, the most effective approaches include the use of self-inactivating KamiCas9 system for editing the huntingtin gene (Merienne et al., 2017), injection of *mHTT* transcription-targeting viral vectors (Zeitler et al., 2019) and application of siRNAs, shRNAs or miRNAs for the reduction of *mHTT* mRNA and protein level (see (Kaemmerer and Grondin, 2019) for review).

It is obvious that gene therapy is becoming more problematic at symptomatic stages of the disease. In HD, as well as other neurodegenerative disorders, late-stage pathology may include numerous compensatory mechanisms which would make it increasingly difficult to implement a causal therapy. But substantial therapeutic benefits may still be possible, notably if basic research can reveal a (virtual) bypass in the signal flow connecting the elements of molecular intervention and motor outcome. In the present experiments with HD mice, a rather robust motor response was induced by local (striatum), cell-selective (astrocytes) and site-directed (EAAT2 C-terminus) transgene expression. Such result is to some extent surprising because most evidence-based models describe the initiation of voluntary movements as the result of a multi-level parallel and distributed information processing in a large number of brain structures (see for instance (Morita, 2014)). That a particular cell population can effectively control a set of motor acts has mostly been implicated in “lower” vertebrates or invertebrates.

However, reports on specific links between motor performance and specific transgene expression are already accumulating (see, for instance, (Nagai et al., 2019;Percie du et al., 2020)) and may help to identify new therapeutic targets. The present results have promise with regard to small molecule therapy of neurologic conditions with hypokinesia, including HD, Parkinson’s disease, stroke and toxin-induced brain damage. These disorders may share similar alterations in the regulation of synaptic glutamate transport, but more information is needed for precise molecular targeting of the involved signaling cascades. The results of bidirectional testing for disease- and treatment-related effects and the combined use of new behavioral and cellular readouts should be a useful complement to the already existing studies on mouse models with advanced HD.

### Which cellular markers report failure or rescue of glutamate clearance?

In the past glutamate uptake has mostly been quantified on the basis of tritium-labeled striatal tissue or synaptosomal preparations (see Introduction). In comparison with the measurement of tissue glutamate uptake, SBFI imaging of individual fluorescence-tagged astrocytes offers the advantage that the uptake activity can directly be compared in transduced *vs.* non-transduced astrocytes. In this way the results should be less affected by other HD-induced alterations, such as changes in the size and density of astroglia in the striatum. Our present analysis provides the still missing proof that the deficits in locomotion and synaptic glutamate clearance were in fact associated with: i) astrocytes expressing an HD phenotype, ii) the EAAT2/GLT-1 transporter and iii) signals mediated via the EAAT2-C-terminal, as opposed to changes in other cell types, transporter proteins or EAAT2 domains.

Unfortunately, the subcellular resolution of glutamate uptake with sodium imaging is not very high, even in the case of focal glutamate uncaging (Dvorzhak et al., 2016). This limits the use of SBFI imaging for the estimation of the glutamate clearance at the sites where it matters - the tripartite synapse. Previous attempts to reveal HD-related clearance deficits with glutamate sensors (Parsons et al., 2016;Parievsky et al., 2017) also failed, most likely due to methodical limitations (slow glutamate sensors, nonselective general tissue depolarization, low temporal and spatial resolution of synaptic responses). Considering, however, the importance of the corticostriatal pathway for the initiation of self-induced movements (Plotkin and Surmeier, 2015) it was found worthwhile to establish a new approach for the evaluation of the glutamate transients at individual glutamatergic synapses in the dorsal striatum (Dvorzhak and Grantyn, 2019). The most telling indicators of clearance deficiency were the time constant of decay and the spread of the stimulus-induced glutamate elevation as reported by the ultrafast glutamate sensor iGlu*u* (Helassa et al., 2018;Dvorzhak et al., 2019). It was already known that the duration of corticostriatal EPSCs increased (Dvorzhak et al., 2019) or, respectively, decreased (Lee et al., 2008;Ghosh et al., 2016) after the exposure to pharmacological blockers of glutamate transport or modulators of NF-κB-dependent transcription. A faster decay of the glutamate elevation at corticostriatal terminals located on S506X-expressing astrocytes can therefore be regarded as a convincing argument in support of the astrocytic contribution to the glutamate clearance in HD. According to previous mathematical modeling (Zheng et al., 2008;Scimemi and Beato, 2009;Kessler, 2013) this could diminish the potentially adverse effect of extrasynaptic NMDA receptor activation.

### What could be the molecular basis of failing glutamate uptake in HD?

In principle, insufficient glutamate uptake could be due to aberrant RNA splicing (Lin et al., 1998;Scott et al., 2011) or changes in the relative abundance of functional EAAT2 isoforms (Goursaud et al., 2011). At least 3 of the latter are known to differ in their C-terminal sequence: EAAT2a, EAAT2b and EAAT2c (GLT-1a, GLT-1b and GLT1c in rodents, see (Vandenberg and Ryan, 2013) for further information). In the rat brain, EAAT2a is ~ 15-fold more abundant than EAAT2b (Holmseth et al., 2009). Both are found at glutamatergic synapses (Al Awabdh et al., 2016), but only EAAT2b is required for the regulated as opposed to constitutive glutamate transport (Underhill et al., 2015). The C-terminus of EAAT2b but not EAAT2a contains a sequence predicted to interact with PDZ-containing proteins, including PICK1 (Wheatley et al., 1992;Bassan et al., 2008), PSD95 (Gonzalez-Gonzalez et al., 2009), or DLG1(Underhill et al., 2015)). By interaction with these proteins, EAAT2 can increase its internalization and/or lateral mobility in the astrocyte membrane due to activation of CamKII (Underhill et al., 2015) or high concentration of glutamate (Al Awabdh et al., 2016). It was shown that experimental immobilization of EAAT2 prolongs synaptic currents (Murphy-Royal et al., 2015). Given that the duration of the glutamate transport cycle (~ 12 ms at Schaffer collateral synapses (Bergles and Jahr, 1998)) is relatively long in comparison with the glutamate receptor binding, it was hypothesized that lateral movement of the substrate-transporter complex and a rapid membrane turnover to replenish fresh unbound transporter molecules are necessary for effective glutamate buffering at synaptic sites. To ensure the high abundance of transporter molecules around a glutamate-releasing terminal (Lehre and Danbolt, 1998) the turnover of the transporter from the cytoplasm into the membrane should also be high. However, how the degradation/replenishment of EAAT2 protein at a tripartite synapse actually works is not yet clear.

The present experiments in symptomatic Q175 HET are, to the best of our knowledge, the first attempt to characterize an HD phenotype in the EAAT2 interactome by using immunoprecipitation of mYFP-tagged native and C-terminal-truncated EAAT2 as “bait”. The focus was on alterations produced by the disease, on one side, and changes due to C-terminal truncation of EAAT2, on the other side. It became quite obvious that HD produced major changes in the abundance of full-length EAAT2 binding partners, while expression of mYFP-EAAT2-S506X resulted in a partial return to the interaction pattern seen in WT.

The changes in HD mice included the disappearance of normally existing, potentially necessary interactors, such as dystrophin, and the appearance of new, potentially pathological interactors, such as UBA2. Dystrophin is quite plentiful in astrocytes, especially at the endfeet, and it has been shown that dystrophin-glycoprotein complexes provide a link between laminin and the cytoskeleton (Sato et al., 2018) thereby contributing to the stabilization of aquaporin4 and Kir4.1 in particular subregions of the astrocyte plasma membrane (Enger et al., 2012). The function of dystrophin at glutamatergic corticostriatal synapses has not yet been explored, but there is evidence for a role of a dystrophin-associated protein complex in the pathogenesis of parkinsonian symptoms (Spuler et al., 2010). UBA2 and SAE1 form a heterodimer that functions as a SUMO-activating enzyme. Sumoylation - the covalent attachment of SUMO1 to lysine residues - is a posttranslational modification process with relevance to HD (Steffan et al., 2004). Interestingly, non-sumoylated EAAT2 resides in the plasma membrane while the product of SUMO fusion with EAAT2 tends to form aggregates in the cytoplasm (Foran et al., 2011). In the present study, when comparing the UBA2 levels precipitated by mYFP-EAAT2 or mYFP-EAAT2-S506X from HET animals, it was apparent that the loss of C-terminal interaction motives largely prevented the interaction between EAAT2 and UBA2. In cultured astrocytes, a proteolytically released C-terminal fragment of EAAT2 modified by SUMO1 was also found in the nucleus, with inhibitory effects on EAAT2 transcription (Gibb et al., 2007;Foran et al., 2011). The relevance of this EAAT2-derived signaling mechanism was demonstrated *in vivo,* in a mutant SOD1 mouse model of amyotrophic lateral sclerosis (ALS),where knock-in of a modified *Slc1a2* isoform with a defective caspase-3 cleavage site prolonged the life span of mice afflicted by the disease (Rosenblum et al., 2017).

As for the plasma membrane localization of EAAT2, it should be noted that there is a basal constitutive internalization of the transporter, with a critical role of the lysines 497, 517, 526, 550, 558, 570, and 573 at the EAAT2 C-terminal (Gonzalez-Gonzalez et al., 2008;Martinez-Villarreal et al., 2012). It is interesting that in the present SBFI imaging experiments a selective lysine-to-arginine exchange in the C-terminal of the EAAT2 protein proved to be almost as effective as the C-terminal truncation suggesting a critical role for C-terminal lysine-mediated interactions such as sumoylation and ubiquitination in the regulation of glutamate uptake.

In the context of the above literature on constitutive internalization of EAAT2 and the toxic effects of sumoylated C-terminal EAAT2 cleavage products, it is tempting to propose that an astrocyte expressing *mHTT* presents with largely altered conditions for both transcriptional control and protein sorting, which perhaps explains why in Q175 HET a mere stimulation of full-length EAAT2 failed to eliminate the deficits in the motor performance and glutamate uptake seen in HET CTRL group. Of course, much more work is needed to fully unravel the molecular determinants of EAAT2 function in the striatum and other regions of the brain.

## METHODS

### AAV plasmid design and vector production

Fig. 1 and Supplemental Tab. 3 present an overview of the used vectors, the sites of injections and the purpose of a chosen indicator. Plasmid pRcCMV-mYFP-EAAT2 and pRcCMV-mYFP-EAAT2-S506X were gifts from Christoph Fahlke and Arnd Baumann (Forschungszentrum Jülich). pRcCMV-mYFP-EAAT2 contains the expression cassette for the human *Slc1a2* fused to N-terminal mYFP. In pRcCMV-mYFP-EAAT2-S506X a serine at position 506 is point-mutated to generate a stop codon that causes the truncation of the last 68 amino acids of the C-terminus. This truncation does not interfere with the membrane insertion of the protein or glutamate transport (Leinenweber et al., 2011). In contrast, the mutation of four C-terminal lysine (Lys, K) residues to arginine (Arg, R) in the full-length EAAT2 has been shown to be critical for transporter internalization (Gonzalez-Gonzalez et al., 2008). Four consecutive cycles of site-directed mutagenesis were applied to generate EAAT2-4KR. In brief, pRcCMV-mYFP-EAAT2 was amplified by PCR using the mutagenesis primers listed in Supplemental Tab. 1, and the template was subsequently destroyed by DpnI digest. For high-efficiency transformation, NEB C2987H^®^ alpha-competent E. coli were incubated with the DpnI-digested PCR mix. Successful mutants were screened by Sanger sequencing (Eurofins Genomics, Köln). The expression cassettes for mYFP-EAAT2, mYFP-EAAT2-4KR and mYFP-EAAT2-S506X were subcloned by restriction digest with BamHI and EcoRI and ligation into a preexisting AAV backbone (pAAV-gfaABC1D-ChR2(LCTC)-p2a-FP635-WPRE) to produce pAAV-gfaABC1D-mYFP-EAAT2-WPRE, pAAV-gfaABC1D-mYFP-EAAT2-4KR-WPRE and pAAV-gfaABC1D-mYFP-EAAT2-S506X-WPRE. Monomeric mRuby3 was subcloned from pKanCMV-mRuby3-10aa-H2B into the three AAV vectors with AgeI and BsrGI to facilitate detection of the transduced astrocytes and the EGFP-based iGlu*u* in the corticostriatal terminals of the same preparation. Control vectors were constructed by amplifying mRuby3 or mYFP by PCR adding a 3’ stop codon and a SalI restriction site and were then cloned into an AAV backbone to generate pAAV-gfaABC1D-mYFP-WPRE or pAAV-gfaABC1D-mRuby3-WPRE. The plasmids used or modified for further use are listed in Supplemental Tab. 1. The AAV vectors were produced at the Vector Core Facility of the Charité - University Medicine or at UPenn Vector Core. We acknowledge the contribution of Viviana Gradinaru and Benjamin Devermann for generating the PHP.eB adeno-associated virus (AAV) serotype (Chan et al., 2017).

### Antibodies

The primary and secondary antibodies used for the quantification of WB or IR are also listed in Supplemental Tab. 3. For concentrations see the Methods section “Quantitative immunofluorescence (IF)” and “ Western blot analysis of striatal lysates”. The EAAT2-Ab (Abcam, Cambridge, UK) was raised against a synthetic peptide within rat EAAT2 aa 550 to the C-terminus (C terminal) conjugated to keyhole limpet haemocyanin. The exact sequence is proprietary. Apart from several other C-terminus-directed Abs, we also tried Ab77039 and Ab203130 from Abcam were peptide fragments between aa143 and 239 were used as antigen but did not achieve the quality of immunostaining seen with Ab41621.

### Animals

Z-Q175-KI mice were obtained from CHDI (“Cure Huntington’s Disease Initiative”, see stock # 027410 of the Jackson Laboratory, Bar Harbor, USA). All applicable international, national and institutional guidelines for the care and use of the animals were followed. The study design, performance of experiments and statistical evaluation has been approved by the Berlin Office of Health Protection and Technical Safety (G0218/17), with a yearly update of the experimental guidelines by the local authorities according to the 10 essential rules of ARRIVE (see latest update from July 14, 2020 and (Percie du et al., 2020)). The experiments were performed in animals of either sex at an age of 51 to 76 weeks. The number of CAG repeats ranged from 182 to 201 and were determined by Laragen (Culver City, CA, USA). Care was taken that all experimental groups contained an equal number of males and females (±1). Apart from the body weight, no systematic differences could be detected in any of the evaluated indicators. Blinding was not applied, since the animals were used at a stage of HD when the experimenters could recognize the respective genotype and select the appropriate mice among a usually very small number of available animals. The necessary sample size was kept to a minimum and was calculated beforehand to achieve hypothesis testing at a significance level of 5%.

### Stereotaxic and intravenous injections of viral expression vectors for treatment and diagnostic purposes

Q175 WT and HET were anaesthetized by intraperitoneal injection of a mixture containing 87.5 mg/kg ketamine and 12.5 mg/kg xylazine (both from Sigma-Aldrich, Taufkirchen) before receiving bilateral intrastriatal injections of PHP.eB-mRuby3-gfaABS1D-EAAT2 (or respective isoforms/controls) as part of the treatment. The therapeutic vectors or controls were given at a concentration of 1.0*10^12^ gc/ml and an amount of 1 μl per site (1.0*10^9^gc/striatum) at the following coordinates with respect to bregma (mm): anterior 0.75, lateral 2.0, ventral 2.5. The intravenous injections (100 μl) were performed under slight isoflurane anaesthesia and adjusted to an injected amount of 2.0*10^11^ to 3.5*10^11^ gc/animal. The expression vector for the glutamate sensor AAV9-CamKII.iGlu*u*.WPRE-hGH (7.34*10^13^gc/ml - 0.3 μl) was applied at four sites at: anterior 1.5, lateral 1.56, 1.8, 2.04, 2.28 and ventral 1.7. The expression time between injection and sacrifice was 3-5 weeks.

### Behavioral tests in WT and Q175 HET

On the day of sacrifice, the animals were submitted to a classical open field test (53 cm x 33 cm), as described in a previous study with Q175 homozygotes (Rothe et al., 2015). The mice were individually tested between 9 a.m. and 2 p.m. using an open-field box made of gray plastic with 53 cm x 33 cm surface and 19 cm walls, illuminated by a spot of 25 W placed 1 m above the shadow-free box. Monitoring of 5-min sessions was done by a video camera (Logitech C525 Webcam, 15 frames/s). A customary software was designed to quantify not only running activity but also the motor activity at rest. This required the calculation of the centroid of all pixels derived from the black mouse body. To this end, the background frame without mouse was subtracted from frames with mouse using an intensity threshold of mean + 3SD. A virtual mouse radius (v.m.r.) was derived from the area of suprathreshold mouse pixels approximated as a circle. The identification of the resting state is based on the time needed to cross 2 v.m.r. For more details and definitions see Supplemental text.

For the “step-over test” the animal was placed into the center of a Petri dish (diameter 18.5 cm and wall height 2.8 cm). The movements were recorded with a video camera. Using offline analysis, a software routine determined the time between the take-off of the experimenter’s hand (in a black glove) and the moment when the animal has reached the outside of the dish with all 4 feet. This parameter was called “step-over latency” and correlated with the open field total path in 5 min.

### Preparation of acute brain slices

The animals were anaesthetized with isoflurane, transcardially perfused with cooled aerated saline containing (in mM): NMDG - 92, KCl - 2.5, NaH2PO4 - 1.25, NaHCO3 - 25, glucose - 20, CaCl_2_ - 0.5, MgCl_2_ - 10, sodium pyruvate - 3, and sodium ascorbate - 5 (pH 7.35, 303 mosmol/l). After decapitation and removal of the brains, parasagittal (10 deg off) sections (300 μm) containing the striatum were prepared as previously described (Dvorzhak et al., 2016). The slices were kept in artificial cerebrospinal fluid (ACSF) containing (in mM): NaCl - 125, KCl - 3, NaH2PO4 - 1.25, NaHCO3 - 25, CaCl2 - 2, MgCl2 - 1, glucose - 10 (pH 7.3, 303 mosmol/l), supplemented with (in mM) sodium pyruvate - 0.5, sodium ascorbate - 2.8 and glutathione - 0.005. These perfusion and recovery solutions preserved the astrocytes better than physiological ACSF, sucrose- or choline-containing solutions, the criterion being the astrocyte resting membrane potential at break-in WT (<= −75 mV).

### Sodium imaging of astrocytes in striatal slices

Our methods for sodium imaging in striatal astrocytes largely followed the techniques already described (Dvorzhak et al., 2016). Briefly, the slices were incubated for 20 - 30 min in oxygenated ACSF containing 222 μM of the membrane-permeable form of SBFI (#S-1264, Thermo Fisher Scientific), and 2.5% dimethyl sulfoxide (DMSO) and 0.5% Pluronic F-127 at 36°C. For recording of the L-aspartate-induced sodium transients, the gap junction blocker CBX (100 μM, Abcam) was added to the superfusion solution, along with blockers of ionotropic glutamate receptor blockers (DNQX 10 μM and MK801 1 μM, Tocris, Bristol, UK). Wide-field fluorescence imaging of SBFI-AM-stained slices was performed using a digital live acquisition imaging system (Andor Solis version 4.30.30034.0, Acal GmbH, Gröbenzell) and a sCMOS camera (AndorZyla 4.2 plus) attached to an upright Zeiss microscope (Axio Examiner A1, Göttingen). Images were collected with a Zeiss 63x NA 1.0 water immersion plan apochromat objective. Cells were selected based on the resting levels of SBFI and mRuby3 fluorescence excited by a UVICO ultraviolet or visible light source (Rapp OptoElectronic, Hamburg), combined with suitable filter sets from Omega Optical (Brattleboro, VT, USA) attached to a FW 1000 filter wheel (Applied Scientific Instrumentation, Eugene, OR, USA). Single wavelength sodium imaging was performed by excitation of SBFI at 380 nm (sodium-sensitive wavelength). SBFI emission was collected at >510 nm (dichroic mirror XF2002, emission filter XF3086, Omega Optical). Regions of interest (ROIs) with a size of 3.2 μm x 3.2 μm were defined on the cell body. Binning was 8 x 8. The spatial resolution was then 0.8 μm/pixel. The exposure times was set to 150 ms in all experiments. Images were acquired every 3 sec. Custom-written software was used to control image acquisition and the valves operating the superfusion system. After 1 min of baseline recordings (20 images), L-aspartate (1 mM) was applied for 1 min followed by a 3 min washout period. Subsequently, the same routine was repeated in the presence of the glutamate transport blocker TFB-TBOA 2 μM (Tocris). For both traces the fluorescence change was calculated from the average of the baseline fluorescence intensities for each ROI as ΔF/F = (F_SBFI_ - F_SBFI(baseline)_/F_SBFI(baseline)_. The difference between the two traces represents the L-aspartate-induced sodium transient that is mediated by all available glutamate transporters. The response to L-aspartate was verified by student’s t-test comparison between response peak and baseline.

### Glutamate imaging at single corticostriatal synapses

The methods established to image glutamate release from single corticostriatal presynaptic terminals and to identify synapses exhibiting an HD phenotype have already been described in some detail (Dvorzhak et al., 2019). Briefly, acute slices were submerged into a perfusion chamber with a constant flow of oxygenated ACSF at a rate of 1-2 ml/min. The temperature during the recordings was maintained at 26 - 27 °C. Single varicosities expressing the ultrafast glutamate sensor iGlu*u* (Helassa et al., 2018) were visualized using a Zeiss wide field microscope (Axioscope 2, FS Plus) with a 63x /NA 1.0 water immersion objective and brief (180 ms) discontinuous exposure to a 473 nm laser beam focused to a circular area of ~4.5 μm in diameter. For evaluation of evoked responses, the iGlu*u* fluorescence was acquired at a frequency of 2.5 kHz from a rectangular ROI of 4 μm x 4 μm (20 x 20 pixels, binning 2) using a sCMOS camera (Andor Zyla4.2 plus). Laser, camera and electrical stimulation of the axon/bouton were controlled by in-house written software routines. Each pixel of the ROI was evaluated separately. The iGlu*u* pixel signal was expressed as change of fluorescence intensity ΔF in % of baseline fluorescence of the given pixel. The baseline is the mean of the intensity values obtained during the 50 ms period prior to stimulation. For the construction of time- and space-dependent [Glu] profiles after evoked release, suprathreshold pixels were determined, the threshold being defined as 3 SD of the ΔF/F baseline. The stimulus-induced changes of suprathreshold ΔF/F in time or space will be referred to as “iGlu_*u*_ transients” or simply “transients”. The term “Peak amplitude” refers to the peak ΔF/F value of an averaged intensity transient derived from all suprathreshold pixels. “Tau decay” or “TauD” is the time constant of decay derived by fitting a monoexponential function to the decay from the peak of the averaged transients. The spatial extension of the iGlu*u* signal is described on the basis of a virtual diameter derived from the area of all suprathreshold pixels combined to form a virtual circle. The area of suprathreshold pixels and the resulting virtual diameter were used as indicators of “Bouton size” (at rest, before stimulation) or “Spread” (after stimulation and glutamate release). The term “Peak spread” refers to the peak value of the averaged spread transient. Dysfunctional synapses could best be detected by analysis of single-pixel iGlu*u*, using the pixel with the highest iGlu*u* elevation at any given terminal. The highest iGlu*u* elevations were always found within or next to the bouton at rest. The peak amplitude of the single pixel transient with the highest iGlu*u* elevation are referred to as “Maximal amplitude”. The respective TauD values are referred to as “TauDmax”.

### Identification of dysfunctional synapses

To induce the glutamate release from individual synaptic boutons under physiological conditions, a depolarizing current pulse was applied through an ACSF-filled glass pipette (tip diameter <1 μM, resistance 10 MΩ) placed next to an axon in close proximity with a fluorescent varicosity. Responses were elicited at minimal intensity at a repetition frequency of 0.1 Hz. To challenge the glutamate uptake mechanisms at the site of release, individual synapses were stimulated under condition of blocked action potential generation (in the presence of tetrodotoxin, TTX, 1 μM and in elevated (5 mM) [Ca^2+^]_ec_. This standardized activation mode bypassed eventually existing disease-related differences in the myelination and excitability of corticostriatal axons. The identification of an HD phenotype in corticostriatal synapses is based on the TauD value of the glutamate transient (Dvorzhak et al., 2019).

### Quantitative immunofluorescence at glutamatergic synapses

Using deep isoflurane anaesthesia, mice were transcardially perfused with 60 ml ice-cold phosphate-buffered saline (PBS) containing 4% (w/v) paraformaldehyde in PBS. Sagittal sections (30 μm) were prepared as previously described (Rothe et al., 2015). For the quantification of the transduction rate and transduction specificity freely floating sections were triple-stained with goat anti-td-Tomato (also detects RFP variants like mRuby3) 1:6000 (Sicgen-Acris, Carcavelos), mouse anti S100β 1:2000 (Novus, Abingdon) and rabbit anti-Iba1 1:1000 (Wako Chemicals GmbH, Neuss) or double-stained with goat anti-td-Tomato and mouse anti-NeuN 1:500 (Merck Millipore, Darmstadt). Counter-staining for nuclei with 42,6-diamidin-2-phenylindol (DAPI) 1:10000 was performed in both experiments. For the quantification of synaptic EAAT2 fluorescence, the sections were triple-stained with goat anti-tdTomato-1:6000 (Sicgen-Acris), guinea pig anti-vGluT1 1:1600 (Synaptic Systems, Göttingen) and rabbit anti-EAAT2 1:2000 (Abcam), followed by respective secondary antibodies, as listed in Supplemental Tab. 1. All sections from the different test groups were stained together for reliable comparison of EAAT2 IF levels.

RGB 24 bit images (1024 x 1024 pixels, pixel size 0.08 μm, 3x zoom, no binning) were acquired from the dorsal striatum using an inverted Leica DMI 6000 confocal microscope with an HCX PL Apo 63x oil objective (NA1.4) and stored in the tiff file format. Areas of interest (AOIs, 400 x 400 pixels) were cropped from the larger viewfields, selecting neuropil areas with a minimum of cell somatas or vessels. Quantification of EAAT2 IF was performed using ImagePro Plus (MediaCybernetics, Roper, Sarasota). For comparison purposes, the same staining conditions and acquisition settings were applied to achieve reliable EAAT2 quantification in 4-5 groups with at least 4 animals per group. Within the selected AOIs, smaller rectangular ROIs (25 x 25 pixels) were centred to individual vGluT1+ spots to determine the level of synaptic EAAT2 IF. A threshold algorithm was used to define the boundaries of the EAAT2+ area excluding pixels with F< ROI mean + 0.5 SD. The data is expressed as integral intensity of suprathreshold pixels. The term “Synaptic integral EAAT2 IF” refers to the mean value from 30 individually assessed ROIs within the boundaries of one transduced (i.e. mRuby3+) astrocyte.

### Preparation of striatal lysates

The animals were anaesthetized with isoflurane, transcardially perfused with cooled aerated saline containing (in mM): N-methyl-D-glucamine chloride (NMDG) - 92, KCl - 2.5, NaH2PO4 - 1.25, NaHCO3 - 25, glucose - 20, CaCl2 - 0.5, MgCl2 - 10, Na pyruvate - 3, and Na ascorbate - 5 (pH 7.35, 303 mosmol/l). If not mentioned otherwise, all chemicals and drugs mentioned here and in the following were obtained from Sigma-Aldrich (Taufkirchen). Brains were quickly removed, placed into a custom-made slicing mould and immersed into ice-cold oxygenated NMDG preparation solution. 2 mm thick sagittal slices where cut at a distance of 1-3 mm from midline. The striata were dissected and snap-frozen in liquid nitrogen. The tissue was pulverized under cryogenic conditions using a cryogrinder set (CG 08-02, OPS Diagnostics, Lebanon, NJ, USA). The samples were solubilized in 200 μl lysis buffer containing (in mM) NaCl 150, Tris pH 7.5 50, n-ethylmaleimide 20, dithiothreitol (DTT) 1 supplemented with glycerol 5%, igepal ca-630 1% and Roche cOmplete protease inhibitor cocktail −1x (all from Sigma-Aldrich), kept on ice for 30 min and homogenized at 5000 rpm using a Polytron PT1300D homogenizer. Cell debris was removed by centrifugation at 14.000 x g for 10 min at 4 °C.

The cleared lysates were either directly submitted to WB analysis or used for the immunoprecipitation of YFP-tagged EAAT2 variants with magnetic beads for subsequent WB analysis or mass spectrometry.

### Western blot analysis of striatal lysates

Samples were prepared according to the NuPAGE Technical Guide of Invitrogen. Briefly, after denaturation in NuPAGE LDS sample buffer with DTT 50 mM for 10 min at 70 °C the samples and markers were run on a Novex bis-tris gradient gel (4-12%, Thermo Fischer Scientific) using NuPAGE MOPS SDS running buffer and subsequently blotted on a Novex 0.45 μm nitrocellulose membrane (LC2001, Thermo Fischer Scientific). The membranes were washed and blocked in ReadyTector solution A (CANDOR Bioscience GmbH, Wangen), and the primary antibodies rabbit anti-EAAT2 1:2000 (Abcam, Cambridge, UK) and mouse anti-GAPDH 1:1000 (Sigma-Aldrich) were directly applied for 1 h at room temperature in solution B that also contained horseradish peroxidase (HRP) coupled to the secondary antibodies against rabbit or mouse, respectively. To detect mYFP, membranes were blocked for 30 min in 1x Roti-block solution (Carl Roth GmbH, Karlsruhe) before applying rabbit anti-GFP 1:1000 (ChromoTek GmbH, Martinsried) over night at 4 °C in the same solution. Then the HRP-coupled secondary antibodies (Dianova GmbH, Hamburg) were applied at 1:2500 for 3 h at 4 °C. Proteins were detected using respective kits from Biozym Scientific GmbH (Hessisch Oldendorf. Proteins were detected by chemiluminescence and submitted to image analysis with ImageJ.

### Immunoprecipitation of YFP-tagged EAAT2 for western blotting or mass spectrometry

Striatal lysates were diluted with 300 μl wash buffer (in mM) NaCl 150, Tris pH 7.5 50, DTT 1, glycerol 5% and incubated for 2 h with 30 μl of pre-washed GFP-Trap magnetic agarose beads (gtma-20, ChromoTek, Planegg-Martinsried) using gentle rotation at 4 °C. The beads-protein complexes were isolated using a DynaMag-2 magnet (Thermo Fisher Scientific), applying 3 wash-resuspension cycles before short-term storage of the immunoprecipitate at −18 °C. The first wash solution contained igepal ca-630 0.05%. For WB analysis the beads were for 10 min incubated in the NuPAGE LDS sample buffer and treated as described in the previous chapter. For liquid chromatography tandem mass spectrometry (LC-MS), the beads were first incubated in digestion buffer (sodium deoxycholate - 1%, dithiothreitol - 10 mM, ammonium bicarbonate - 50 mM, 45 min, r. t.). The proteins were then submitted to alkylation with 55 mM chloroacetamide (30 min, r. t. complete darkness) and over-night digestion with 500 ng endopeptidase LysC (Wako, Neuss) and 500 ng sequence grade trypsin (Promega, Mannheim, GER) at 37°C. The samples were then acidified with formic acid (final concentration 1%).

### EAAT2 interaction proteomics

The peptides were extracted and desalted using the StageTips protocol. Separation was carried out using in-house-manufactured 20 cm fritless silica microcolumns with an inner diameter of 75 μm, packed with ReproSil-Pur C18-AQ 1.9 μm resin (Dr. Maisch GmbH, Ammerbuch), a 98 min gradient with a 250 nl/min flow rate of increasing Buffer B concentration (from 2% to 60%, Buffer B: 90% acetonitrile) on an High Performance Liquid Chromatography (HPLC) system from Thermo Fischer Scientific. The eluting peptides were directly ionized by electrospray ionization and transferred into a Thermo Orbitrap Fusion mass spectrometer. The instrument was operated in the data-dependent mode with performing full scans in Orbitrap (60K resolution; 4 x 10^5^ ion count target; maximum injection time 50 ms), followed by top 20 MS2 scans using higher-energy collision dissociation (NCE of 32; 15K resolution, 5 x 10^4^ ion count target; 0.7 m/z isolation window; maximum injection time: 250 ms). Only precursor with charge states between 2-7 were fragmented. Dynamic exclusion was set to 30 sec. Raw data were analyzed using the MaxQuant software (v1.6.0.1). The internal Andromeda search engine was used to search MS2 spectra against a decoy UniProt database for mouse (MOUSE.2018-05), as well as the sequences of the mYFP fusion constructs, containing forward and reverse sequences. The search included variable modifications of oxidation (M) and N-terminal acetylation, deamidation (N and Q) and fixed modification of carbamidomethyl cysteine. Minimal peptide length was set to 7 amino acids and a maximum of two missed cleavages was allowed. The false discovery rate (FDR) was set to 0.01 for peptide and protein identifications. The integrated label-free quantification and IBAQ calculation algorithm were activated. Unique and razor peptides were considered for quantification. Retention times were recalibrated based on the built-in nonlinear time-rescaling algorithm and MS/MS identifications were transferred between LC-MS/MS runs with the “Match between runs” option, in which the maximal retention time window was set to 0.7 min. The resulting text files were used for further analyses using the Perseus software package (omicX, v. 1.6.2.1). LFQ intensity values were used for quantification. Reverse hits, contaminants and proteins only identified by site were filtered out. Technical and biological replicates for each condition were defined as groups and intensity values were filtered for “minimum value of 3” per group. After log2 transformation missing values were imputed with random noise simulating the detection limit of the mass spectrometer. Imputed values are taken from a log normal distribution with 0.25× the standard deviation of the measured, logarithmized values, down-shifted by 1.8 standard deviations. The data was obtained from 4 test groups, each comprising at least 4 animals per group. The signals obtained from any given animal (pooling the tissue from both striata) were normalized to the mYFP signal median intensity calculated from all samples. Only proteins with >3 peptide detection and abundance of log2 intensities >= 23 in all 4 samples were included. Differential protein abundance was calculated using two-sample Student’s t test, applying a permutation-based FDR cut-off at 1%. Abundance differences between the samples with a p-value of <=0.05 in a two-tailed t-test were considered significant.

### Data evaluation and statistical analysis

The comparison of the means could be influenced by interanimal variance. Therefore, in the case that individual sections, cells or synapses were obtained from different animals, multilevel (“nested data”) analysis was performed with Prism 8 (GraphPad, San Diego, USA). P values of <0.05 were considered statistically significant. Differences between the groups were tested with ANOVA or respective nonparametric methods (Kruskal-Wallis-test), followed by multiple comparison (Dunnet’s or Dunn’s tests). Significance levels were marked by asterisks, where * corresponds to P<0.05, ** - P<0.01 and *** - P<0.001. The HET-CTRL data served as reference for comparison with WT-CTRL or HET-TEST. Effect strength was described according to Cohen’s D or Hedges’ G. D or G values larger 0.8 suggest that the respective effect was strong.

## ACKNOWLEDGEMENTS

Thanks are due to Profs. Arnd Baumann and Christoph Fahlke, ForschungszentrumJülich for the EYFP EAAT2 S506X plasmid, Prof. Francisco Zafra, Universidad Autónoma de Madrid, for important information on the 4KR-edited EAAT2 isoform, Dr. K. Török, St. George’s, University of London, and Dr. N. Helassa, University of Liverpool, for the iGlu*u* plasmid. Prof. Gudrun Ahnert-Hilger, Charité Universitätsmedizin Berlin, and Dr. Hannes Schmidt, Eberhard Karls Universität Tübingen, contributed numerous helpful suggestions. D. Betances and A. Schönherr provided excellent technical assistance. This work was supported by CHDI (A-12467), the German Research Foundation, under Germany’s Excellence Strategy (Exc 2049 - 390688087) and intramural Charité Research Funds to R.G.

