## Supplemental data and text file for "Expression of a modified astrocytic glutamate transporter alleviates Huntington’s hypokinesia, promotes synaptic glutamate clearance and counteracts potentially adverse EAAT2 interactions"

### SUPPLEMENTAL MATERIALS

### 3 TABLES

### TEXT WITH 4 FIGURES

**Supplemental Table 1.** Summary of results.

| HD phenotype | WT-CTRL | | | | HET-CTRL | | | | HET-EAAT2-S506X | | | | Statistics | | | Posthoc MC HET-CTRL vs. HET-S506X | | |
|---|---|---|---|---|---|---|---|---|---|---|---|---|---|---|---|---|---|---|
| | Mean | SE | N-c/t | N-a | Mean | SE | N-c/t | N-a | Mean | SE | N-c/t | N-a | Test | F (DFn, Dfd) | P | Test | P | Hedges' G |
| #CAG repeats | nd | | | | 186.1 | 1.2 | | 22 | 185.6 | 1.7 | | 15 | t-test | t=0.24 (df=35) | ns | | | |
| Body weight at sacrifice | 31.28 | 0.86 | | 21 | 25.35 | 0.77 | | 22 | 26.41 | 1.08 | | 15 | ANOVA | 0.29 (2, 55) | <0.0001 | Tukey | ns | |
| **OF total distance travelled (m/5 min)** | **18.72** | **0.93** | | **14** | **12.99** | **1.77** | | **15** | **19.16** | **1.80** | | **21** | **ANOVA** | **4.09 (2, 47)** | **0.0230** | **Tukey** | **0.027** | **-0.81** |
| **OF #starts/stops (in 5 min)** | **61.07** | **4.78** | | **14** | **36.80** | **6.23** | | **15** | **58.67** | **6.66** | | **21** | **Kruskal-Wallis** | **4.17 (2, 47)** | **0.0216** | **B-K-Y** | **0.040** | **-0.78** |
| **Step-over latency (s)** | **56.7** | **10.00** | | **18** | **179.1** | **46.6** | | **19** | **51.6** | **9.8** | | **14** | **Kruskal-Wallis** | **7.37 (2, 47)** | **0.0251** | **B-K-Y** | **0.041** | **0.79** |
| **Maximal/Mean rest radius** | **2.53** | **0.04** | | **14** | **2.81** | **0.11** | | **15** | **2.58** | **0.21** | | **21** | **ANOVA** | **4.64 (2, 47)** | **0.01** | **Tukey** | **0.038** | **0.75** |
| **Mean running velocity (mm/s)** | **99.53** | **5.14** | | **14** | **81.57** | **6.04** | | **15** | **109.45** | **8.06** | | **21** | **ANOVA** | **4.15 (2, 26)** | **0.0220** | **Tukey** | **0.017** | **-0.87** |
| **Mean velocity at rest (mm/s)** | **37.39** | **1.78** | | **14** | **26.48** | **3.69** | | **15** | **36.91** | **3.12** | | **21** | **ANOVA** | **3.72 (2, 47)** | **0.0318** | **Tukey** | **0.048** | **-0.70** |
| **Open field center time (%)** | **31.48** | **4.32** | | **14** | **14.98** | **3.67** | | **15** | **16.57** | **2.47** | | **21** | **ANOVA** | **6.59 (2, 47)** | **0.0030** | Tukey | ns | |
| **L-Asp-induced Na transient amp (ΔF/F%)*** | **3.30** | **0.26** | **46** | **4** | **2.01** | **0.18** | **36** | **3** | **3.28** | **0.23** | **23** | **3** | **Nested ANOVA** | **4.525 (3 10)** | **0.0299** | **Dunnett** | **0.049** | **-1.0887** |
| **Syn Glu transient(max) spread (μm)** | **1.28** | **0.06** | **35** | **10** | **1.55** | **0.09** | **32** | **16** | **0.94** | **0.08** | **29** | **8** | **Nested ANOVA** | **10.90 (2, 31)** | **0.0003** | **B-K-Y** | **<0.001** | **1.27** |
| **Syn Glu transient(max) TauD (ms)** | **2.76** | **0.29** | **35** | **10** | **9.57** | **1.89** | **32** | **16** | **2.57** | **0.31** | **28** | **8** | **Nested ANOVA** | **5.04 (2, 31)** | **0.0127** | **B-K-Y** | **0.017** | **0.88** |
| Syn Glu transient(max) amp (ΔF/F%) | 85.6 | 11.7 | 35 | 10 | 67.8 | 7.5 | 32 | 16 | 53.5 | 8.3 | 28 | 8 | Nested ANOVA | 0.76 (2,31) | 0.4751 | B-K-Y | ns | 0.33 |
| **Syn EAAT2 IF total (a.u.)*** | **5230** | **168.7** | **12/360** | **2** | **3663** | **81.42** | **12/360** | **2** | **4638** | **205.8** | **12/360** | **2** | **Nested ANOVA** | **12.25 (3,4 )** | **0.0175** | **Dunnett** | **0.048** | **-1.87** |
| **SPN input resistance (MOhm)** | **114** | **10.3** | **18** | **8** | **165** | **17.0** | **14** | **8** | **127** | **13.5** | **8** | **5** | **Nested ANOVA** | **4.05 (2, 37)** | **0.0256** | B-K-Y | 0.104 | 0.64 |
| **ITonic(GABA) amp (pA)** | **11.33** | **2.37** | **13** | **6** | **2.99** | **1.03** | **14** | **8** | **5.78** | **3.37** | **7** | **5** | **Nested ANOVA** | **6.19 (2, 16)** | **0.0102** | B-K-Y | ns | |

*Test includes 4KR group
Abbreviations: MC - multiple comparison test, BKY - Benjamini, Krieger, Yekutieli test

**Supplemental Table 2.** Definitions of Open Field Test variables.

***<u>Mouse centroid position</u>***

It reflects the body positions in the horizontal plane and is defined as

$$X_c = \frac{\sum_{i=1}^{N} X_i \times Pixel\ intensity_i}{\sum_{i=1}^{N} Pixel\ intensity_i}$$

*Xc, Yc is the  position centroid.  N - the number of suprathreshold pixels, i – pixel index. Xi, Yi – pixel coordinates with corresponding pixel intensity.*

***<u>Virtual mouse radius (v.m.r.)</u>***

$$Virtual\ mouse\ radius = \sqrt{\frac{1}{\pi} \times N_{suprat\ hreshold\ pixels} \times Pixel\ area}$$

***<u>Start point</u>***

*Start point = ! $(X_c,Y_c)$ |• $((X_{Stop}-X_c)^2+(Y_{Stop}-Y_c)^2)$>1×Virtual mouse radius*
*where C(Xc, Yc) - is the current centroid position and Xstop, Ystop – the coordinates of the preceding stop point.*

***<u>Stop point</u>***

*Stop point = $C(X_c,Y_c)$|(• $((X_c-X_{break}{}^2+(Y_c-Y_{break}{}^2)$<1×Virtual mouse radius)U(! $(X_c,Y_c)$=! $(X_0,Y_0)$)*
*where $C(X_0, Y_0)$ is the centroid position of the first frame and $X_{break}$, $Y_{break}$ – the coordinates of the mouse centroid at the moment when it lags the current position by the time of running start.*

***<u>The running start episode</u>***

is an initial component of the open field locomotion between a start and a stop point.
*Running start episode = {$C(X_c,Y_c)$õRunning episode|• $((X_c-X_{Stop})^2+(Y_c-Y_{Stop})^2)$>1×Virtual mouse radius∩• $((X_c-X_{Stop})^2+(Y_c-Y_{Stop})^2)$<2×Virtual mouse radius}*
*where Xstop, Ystop are the coordinates of the preceding stop point.*

***<u>The running episode</u>***

is a component of the open field locomotion between a start and a stop point.
*Running episode = {$C(X_c,Y_c)$õTrajectory[Start point, next Stop point)}*

***<u>The resting episode</u>***

is a component of open field locomotion between a stop and a start point.
*Resting episode = {$C(X_c,Y_c)$ õTrajectory[Stop point, Start point)}*

***<u>Mean running velocity</u>***

$$Mean\ running\ velocity = \frac{\sum_{i=1}^{T} Distance\ travelled\ during\ the\ running\ episode_i}{\sum_{i=1}^{T} Duration\ of\ running\ episode_i}$$

*where T - the number of running episodes duringthe 5 min observation period; i - index of the running episode.*

***<u>Mean resting velocity</u>***

$$Resting\ velocity = \frac{\sum_{i=1}^{R} Position\ change\ during\ the\ resting\ episode\ _i}{\sum_{i=1}^{R} Duration\ of\ resting\ episode\ _i}$$

*where R - the number of resting episodes during the 5 min observation period; i - index of the resting episode.*

***The resting area***

is a circular area centered to the origin of a radius equaling the maximal deviation from the center point.

***The center point of the resting area***

$$X_{center} = \frac{\sum_{i=1}^{N} X_i}{N}; Y_{center} = \frac{\sum_{i=1}^{N} Y_i}{N}$$

*where Xcenter and Ycenter are the coordinates of the center point; N – the number of centroid positions contributing to the resting episode; i – index of centroid value at rest; Xi and Yi - actual coordinates during the resting episode.*

***Maximal deviation from the resting area center point***

$$Maximal\ deviation\ from\ center\ point = Max\left(\sqrt{(X_i - X_{center})^2 + (Y_i - Y_{center})^2}\right)$$

***Mean deviation from the resting area center point***

$$Mean\ deviation\ from\ center\ point = Mean\left(\sqrt{(X_i - X_{center})^2 + (Y_i - Y_{center})^2}\right)$$

***Open field resting time***

%Fraction of the observation time occupied by resting episodes

***Open field center time***

%Fraction of the observation time spent in the central area of the open field

### SUPPLEMENTAL TEXT AND FIGURES

#### Assessment of spontaneous locomotor activity in WT and HD mice

The open field test (OFT) is a frequently used but not highly standardized tool to evaluate spontaneous locomotor activity (Wahlsten, 2001;Kostrzewa and Kas, 2014). Originally it was developed for the characterization of behavioral phenotyping of rodents exhibiting symptoms of anxiety. Later on, it was used for the quantification of symptoms and recovery in rodents with experimental lesions, intoxication and neurodegenerative disease (Crawley, 2003), taking advantage of automated analysis of positional data from movement trackers (Tatem et al., 2014). In many cases the OFT provided not much more than the distance travelled during the time of observation, but a relatively recent study in CAG140 knock-in mice extracted 6 indicators characterizing the locomotion in HET of 52 weeks and older (Fowler and Muma, 2015). These indicators were: distance travelled (1), force variability (2), number of wall rears (3), wall rear duration (4), number of low mobility bouts (5) and number of long strait runs (6). Significant deficits were found in 1, 3, 5, 6. The algorithm used to determine the number of "mobility bouts" and number of "long strait runs" is perhaps similar to our measure "incidence of starts" (as the number of starts in 5 min). We also determined the respective movement velocities and the actual extension of the "bouts". The results were correlated with a new test, the step-over latency test (SOLT), a test to some extent corresponding to the number of wall rears (3) in the Fowler and Muma study. Supplemental Tab. 3 presents the definitions of the analyzed OFT variables. More details are described in the Methods section.

The SOLT is associated with, but not identical to, the OFN. It quantifies the time needed to climb the walls of a large Petri dish (see Methods). In principle, it might be understood as the time needed to initiate exploratory behavior from a forced position in the center of a small platform. However, more work is needed to clarify possible contributions of other determinants of spontaneous motor behavior, first of all anxiety. While HD mice spend significantly less time in the center of an open field, they may need more time to escape from such position to the more comfortable peripheral parts of the open field.

As all behavioral data presented in the main text are derived from mice with viral injections into the brain it seemed useful to compare some of the obtained results with data from noninjected mice. Supplemental Fig.1 presents the outcome of SOLT in noninjected WT and HET at an average age of 59 weeks (HET), i.e. about the same age as the injected mice (compare Fig. 2). No significant differences were found in the step-over latencies of CTRL-injected and non-injected WT and HET.

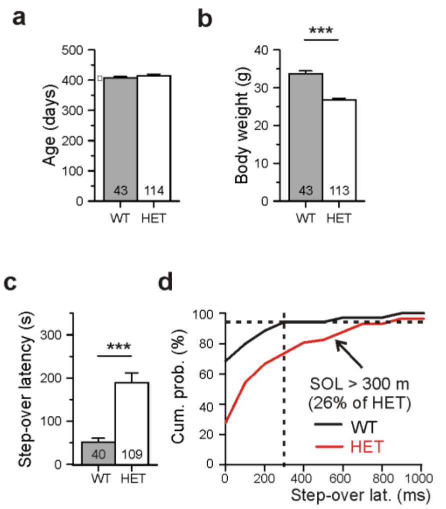


***Supplemental Fig. 1. Results of step-over latency test from noninjected aged WT and Q175 HET.*** *The mean number of CAG repeats in HETs was 191.9 ± 0.76.* ***a-c*** *Mean values of age,body weight and step-over latency.* ***d*** *Cumulative probability plot for step-over latencies of noninjected WT and HET. Note that step-over latencies >300 ms were almost absent in WT, but found in 26% of HET tested once before the viral injection.*

**Failure of full-length EAAT2 to alleviate hypokinesia and to rescue glutamate uptake after intrastiatal injections**

As an additional control for the results presented in Figs. 2 and 3, a construct encoding full-length EAAT2 was injected in the dorsal striatum. Supplemental Fig. 2 presents the respective quantification indicating that Q175 HET injected with the full-length EAAT2 vector failed to benefit from the transgene expression. Likewise, there was no recovery in the glutamate uptake activity of mRuby-positive astrocytes.

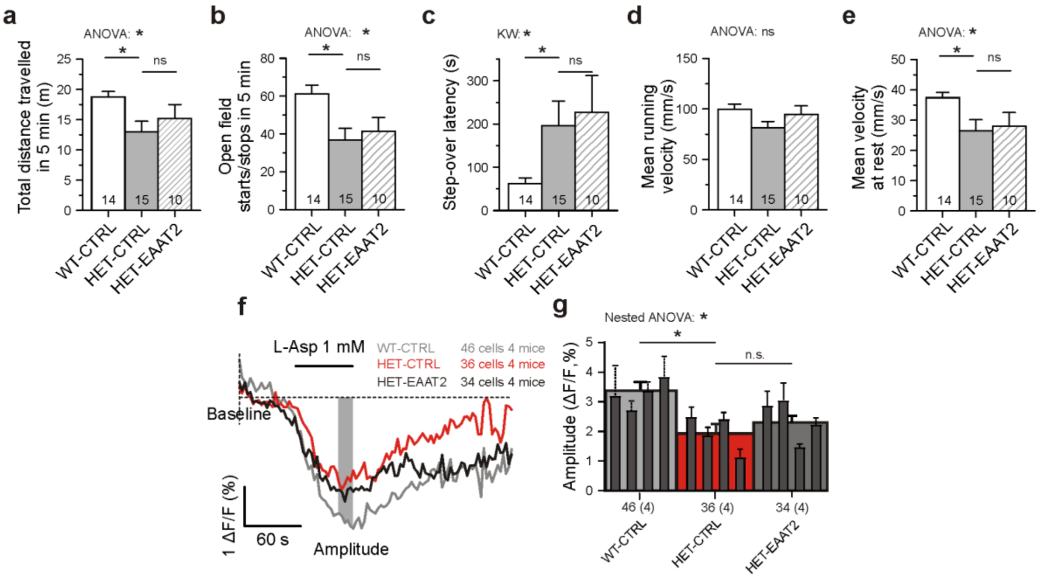


***Supplemental Fig. 2. Failure of intrastriatal full-length EAAT2 to alleviate the motor symptoms of HD and to rescue astrocytic glutamate uptake.*** *Bilateral striatal injections of a PHP.eB-EAAT2 vector with a gfaABC1D promoter and mRuby3 as fluorescent tag. The injected animals were matched for age, body-weight and the number of CAG-repeats.* ***a-e*** *There was no recovery from the motor signs of HD (post-hoc comparison of HET- CTRL vs. HET-EAAT2).* ***f*** *Measurement of single-cell glutamate uptake by sodium imaging with SBFI. Records from mRuby3-positive cells only, in the presence of CBX (100 µM), DNQX (10 µM) and MK801 (1 µM). The sodium response to bath-applied L-aspartate (L-Asp) was defined as the difference between the traces recorded in the absence and presence of TFB-TBOA (2 µM). The amplitude of this differential response was expressed as ΔF/F and derived from the data points during the last 15 sec of L-Asp application. F is the mean fluorescence at rest, before drug application. Averaged traces from all mRuby3+ astrocytes of a group (4 animals per group, 3 groups tested).* ***g*** *Quantification of results with two-level ("nested") ANOVA statistics (animal level and cell level) and two-level post-hoc tests for the differences between the test groups. Here and in all other figures: n.s. - not significant. *, **, *** - p values <.05, 0.01 and 0.001, respectively.*

**Rescue of neuron excitability and astroglial GABA release**

It is already well known that HD progression is associated with an increase in the excitability of striatal projection neurons of the dorsal striatum (SPNs), and it was shown that nonsynaptic release of GABA from astrocytes can contribute to the underlying pathogenic mechanism. The enhanced excitability of striatal neurons is reflected in the appearance of pathological activity patterns (Rothe et al., 2015), lower firing thresholds of individual SPNs and higher values of their whole-cell input resistance $R_N$ (Dvorzhak et al., 2013). Previous studies from our lab suggested that nonsynaptic release of GABA from *mhtt*-expressing astrocytes could contribute to these changes (Wojtowicz et al., 2013).

A similar approach was now used to record the membrane currents of SPNs surrounded by YFP-tagged astrocytes to determine whether changes in $R_N$ could contribute to the phenotype of aged Q175 HET. This was the case (Supplemental Fig. 3a, b and Supplemental Tab. 1). Moreover, EAAT2-S506X-treated HETs exhibited a tendency to change towards WT levels, although a larger number of tests were needed to confirm this possibility with a respective posthoc test.

Patch clamp recordings were further performed to record the tonic GABA(A) receptor currents of SPNs, $I_{Tonic(GABA)}$ (Supplemental Fig. 3c). This current reflects the ambient GABA concentration and, in part, depends on spontaneous release of GABA from astrocytes. The level of extrasynaptic GABA-dependent chloride conductance is believed to play a major role in the control of neuron excitability (Kersante et al., 2013). Due to the co-localization of EAAT2 and GAT3 at the sites of glutamate release and the fact that glutamate transport via EAAT2 can influence the driving forces for GAT3 (Heja et al., 2019;Dvorzhak et al., 2013), a weakness of EAAT2 may lead to a reduction of astrocytic

GABA release via GAT3 and, hence, smaller tonic inhibition of SPNs (Wojtowicz et al., 2013). The present results (Supplemental Fig. 3d, Supplemental Tab. 1) are consistent with this hypothesis although the difference between CTRL- and EAAT2-S506X-treated HET failed to reach the level of statistical significance with nested ANOVA.

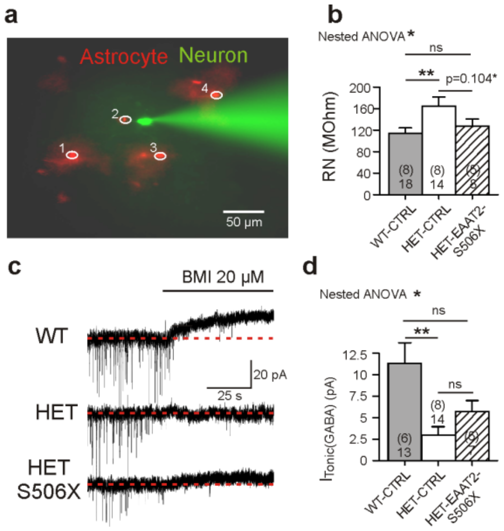


*Supplemental Fig. 3.* ***Neuron excitability and tonic GABA release.*** *Experiments with three test groups ("WT-CTRL", "HET-CTRL" and "HET-EAAT2-S506X"). The expression time of the age- and body-weight-matched mice was 3 weeks. The mean number of CAG repeats showed no difference between the CTRL and treated HET.* ***a*** *Experimental arrangement. SPNs (bright cell with electrode in the center) surrounded by 3-4 astrocytes (in red) were selected for patch clamp recordings, and small hyperpolarizing pulses were delivered through the somatic electrode for the estimation of* $R_N$. ***b*** *Quantification of* $R_N$. ***c, d*** *Traces of* $I_{Tonic(GABA)}$ *and its quantification. Records in the presence of CBX, DNQX and MK801. The numbers in brackets are animals, the numbers below - number of tested SPNs. The differences in c and d were significant (*$p<0.05$*) with two-level "nested" ANOVA statistics. Horizontal bars refer to the post-hoc comparison of the group means.*

**Intravenous injection of EAAT2-S506X failed to produce a significant recovery from hypokinesia although transduced (mRuby+) striatal astrocytes exhibited a significant increase of L-aspartate-induced glutamate uptake**

By chosing intrastriatal injections of the EAAT2 vectors one naturally neglects astrocyte pathologies in other brain areas, including those of the subthalamic nucleus (Atherton et al., 2016), a target of the hyperdirect pathway. Quite likely, one can expect additional or even different effects after systemic application of a given vector.

The tested animal groups were the same as in Fig. 2, 3 (intrastriatal injections) but now EAAT2-S506X and the respective CTRL vectors were injected systemically, via the tail vein. The results are shown in Supplemental Fig. 4. None of the quantified locomotor indicators showed a difference between the HET-CTRL and the HET-S506X group. Moreover, the WT CTRL values obtained in the OFT test (both total distance travelled and center time) were significantly reduced in comparison with respective values from WT-CTRL injected in the striatum. These results hinted the possibility that the motor activity was somewhat compromised even in WT-CTRL suggesting that the systemic vector injection might have some side effects. Nevertheless, the tested mRuby+ astrocytes exhibited a partial rescue of the L-aspartate-induced $Na^+$ elevation. It should be mentioned that the label intensity and the number of labeled cells encountered in any given slice were smaller than in mice with striatal injections. Only the brightest astrocytes were selected for the SBFI imaging. The results suggest that the intravenously injected TEST vector EAAT2-S506X was indeed able to increase glutamate uptake activity although the abundance of the transgene appeared to be rather low after intravenous injection.

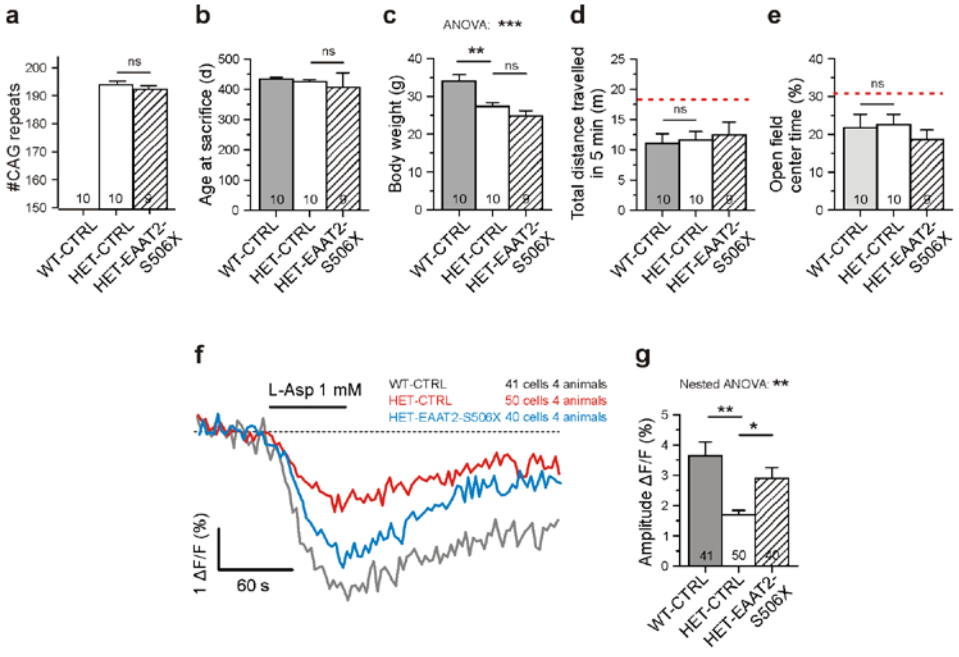


***Supplemental Fig. 4. Lack of behavioral recovery after intravenous injection of EAAT2-S506X., despite partial recovery of astrocytic glutamate uptake in transduced astrocytes. a-c*** *Basic characteristics of tested animals in the experiments with intravenous injection of PHP.eB-gfaABC1D-mRuby3 (3.7E11 gc/animal) in WT and PHP.eB-gfaABC1D-mRuby3-EAAT2-S506X in HET in HET (2.7E11 gc/animal).* ***d, e*** *Results of open field test. Note that in the CTRL-injected WT the performance was worse than in mice injected with the same construct in the striatum. The dotted lines correspond to the respective mean values in Fig. 4d, e.* ***f*** *Averaged traces from the same three test groups. Recording conditions as in Figs. 2 and 4.* ***g*** *Quantification of results.*

Together, these additional control experiments underline the beneficial effects of intrastriatal injection of a C-terminal-modified EAAT2 construct.

**Supplemental Table 3.** Key resources.

| RESOURCE | SOURCE | IDENTIFIER |
|---|---|---|
| ***Animals*** | | |
| Z-Q175-KI | Jackson Labs, Maine, USA | Stock# 027410 |
| ***Antibodies*** | | |
| Donkey anti-Goat IgG, Alexa Fluor® 555 (1:800) | Life Technologies, Darmstadt, GER | Catalogue #A-21432 RRID: AB_2535853 |
| Donkey anti-Guinea pig IgG, Cy™5 (1:200) | Jackson ImmunoResearch, Biozol, Eching, GER | Catalogue #706-175-148 RRID: AB_2340462 |
| Donkey anti-Mouse IgG, Alexa Fluor® 488 (1:800) | Life Technologies, Darmstadt, GER | Catalogue #A-21202 RRID: AB_141607 |
| Donkey anti-Rabbit IgG, Alexa Fluor® 488 (1:800) | Life Technologies, Darmstadt, GER | Catalogue #A-21206 RRID: AB_141708 |
| Donkey anti-Rabbit IgG, Alexa Fluor® 647 (1:200) | Life Technologies, Darmstadt, GER | Catalogue #A-31573 RRID: AB_2536183 |
| Goat anti-tdTomato (1:6000) | Origene Europe, Herford, GER | Catalogue #AB8181-200 RRID: AB_2722750 |
| Guinea pig anti VGluT1 (1:1600) | Synaptic Systems, Göttingen, GER | Catalogue #135 304 RRID: AB_887878 |
| Mouse anti S100B Antibody (SA-12) (1:2000) | Novus, Abingdon, UK | Catalogue #NBP1-41373 RRID: AB_2184561 |
| Mouse anti-NeuN (1:500) | Merck Millipore, Darmstadt, GER | Catalogue #MAB377 RRID: AB_2298772 |
| Rabbit anti EAAT2 (1:2000) | Abcam, Cambridge, UK | Catalogue #ab41621 RRID: AB_941782 |
| Rabbit anti Iba1 (1:1000) | Wako Chemicals GmbH, Neuss, GER | Catalogue #019-19741 RRID: AB_839504 |
| ***Plasmids*** | | |
| pAAV-gfaABC1D-mRuby3 | This study | |
| pAAV-gfaABC1D-mRuby3 EAAT2 | This study | |
| pAAV-gfaABC1D-mRuby3 EAAT2 4KR | This study | |
| pAAV-gfaABC1D-mRuby3 EAAT2 S506X | This study | |
| pAAV-gfaABC1D-mYFP | This study | |
| pAAV-gfaABC1D-mYFP EAAT2 | This study | |
| pAAV-gfaABC1D-mYFP EAAT2 S506X | This study | |
| pCI-syn-iGluu | Addgene, Watertown, MA, USA | Catalogue #106122 |
| pKanCMV-mRuby3-10aa-H2B | Addgene, Watertown, MA, USA | Catalogue #74258 |
| pRcCMV mYFP EAAT2 (K320) | Gifts from C. Fahlke and A.Baumann, Jülich, GER | |
| pRcCMV mYFP EAAT2 S506X (K103) | Gifts from C. Fahlke and A.Baumann, Jülich, GER | |
| ***Primers for site-directed mutagenesis*** | | |
| GLT1-K517R-Fw | gatattgaaatgaccaGgactcaatccatttatg | Modified from González-González et al. 2008 |
| GLT1-K517R-Rw | cataaatggattgagtcCtggtcatttcaatatc | Modified from González-González et al. 2008 |
| GLT1-K526R-Fw | catttatgatgacatgaGgaaccacagggaaag | Modified from González-González et al. 2008 |
| GLT1-K526R-Rw | ctttccctgtggttcCtcatgtcatcataaatg | Modified from González-González et al. 2008 |
| GLT1-K550R-Fw | catagtagatgaatgcaGggtaactctggcag | Modified from González-González et al. 2008 |
| GLT1-K550R-Rw | ctgccagagttaccCtgcattcatctactatg | Modified from González-González et al. 2008 |
| GLT1-K570R-Fw | gaggaagaaccttggaGacgtgagaaataag | Modified from González-González et al. 2008 |
| GLT1-K570R-Rw | cttatttctcacgtCtccaaggttcttcctc | Modified from González-González et al. 2008 |
| ***Recombinant virus strains*** | | |
| pAAV PHP.eB-gfaABC1D-mRuby3 | Charité - Viral Core Facility, Berlin, GER | |
| pAAV PHP.eB-gfaABC1D-mRuby3 EAAT2 | Charité - Viral Core Facility, Berlin, GER | |
| pAAV PHP.eB-gfaABC1D-mRuby3 EAAT2 4KR | Charité - Viral Core Facility, Berlin, GER | |
| pAAV PHP.eB-gfaABC1D-mRuby3 EAAT2 S506X | Charité - Viral Core Facility, Berlin, GER | |
| pAAV PHP.eB-gfaABC1D-mYFP | Charité - Viral Core Facility, Berlin, GER | |
| pAAV PHP.eB-gfaABC1D-mYFP EAAT2 | Charité - Viral Core Facility, Berlin, GER | |
| AAV9-CamKII-iGluSnFRf-u #38698 | UPenn Vector core, Pennsylvania, USA | |
| ***Chemicals, Reagents*** | | |
| 10x Roti-block solution | Carl Roth GmbH, Karlsruhe, GER | Catalogue #A151.1 |
| APV | Abcam, Cambridge, UK | Catalogue #ab120271 |
| Bicuculline methodide | Sigma-Aldrich, Taufkirchen, GER | Catalogue #14343-50MG |

| | | |
|---|---|---|
| Carbenoxolone | Abcam, Cambridge, UK | Catalogue #ab143590-5g |
| DNQX | Tocris, Bristol, UK | Catalogue #2312/50 |
| Endopeptidase LysC | Wako, Neuss, GER | |
| GFP-trap magnetic agarose beads | ChromoTek, Planegg-Martinsried, GER | Catalogue #gtma-20 |
| Ketamine | Sigma-Aldrich, Taufkirchen, GER | Catalogue #K2753-1G |
| MK801 | Sigma-Aldrich, Taufkirchen, GER | Catalogue #M107-50mg |
| Novex 0.45 µm nitrocellulose membrane | ThermoFischer Scientific, Dreieich, GER | Catalogue #LC2001 |
| Novex bis-tris gradient gel 4-12% | Abcam, Cambridge, UK | Catalogue #NP0321BOX |
| Pluronic F-127 | Sigma-Aldrich, Taufkirchen, GER | Catalogue #P2443-250G |
| Ready Tector Solution A | Candor Bioscience G,bH, Wangen, GER | |
| ReadyTector | CANDOR Bioscience GmbH, Wangen, GER | Catalogue #730 040 |
| Roche cOmplete protease inhibitor cocktail | Sigma-Aldrich, Taufkirchen, GER | Catalogue #4693132001 |
| SBFI-AM | ThermoFischer Scientific, Dreieich, GER | Catalogue #S-1264 |
| Sequencing grade trypsin | Promega, Mannheim, GER | Catalogue #V511A |
| Tetrodotoxin | Abcam, Cambridge, UK | Catalogue #ab120054 |
| TFB-TBOA | Tocris, Bristol, UK | Catalogue #2532/10 |
| Trypsin | Promega, Mannheim, GER | |
| WesternBright ECL | Biozym Scientific , Hessisch Oldendorf, GER | Catalogue #R-03031-D25 / R-03025-D25 |
| Xylazine | Sigma-Aldrich, Taufkirchen, GER | Catalogue #110677 |
| ***Equipment*** | | |
| Andor Zyla4.2 plus | Oxford Instruments, Oxford, UK | |
| Axio Examiner A1 | Carl Zeiss AG, Oberkochen, GER | |
| Axioscope 2 FS plus | Carl Zeiss AG, Oberkochen, GER | |
| Cryo-grinder set | OPS Diagnostics, Lebanon, NJ, USA | Catalogue #CG 08-02 |
| DynaMag-2 magnet | ThermoFischer Scientific, Dreieich, GER | |
| Emission filter XF3086 | Omega Optical, Brattleboro, VT, USA | |
| EPC-8 | List, Darmstadt, GER | |
| FW 1000 filter wheel | Applied Scientific Instrumentation, Eugene, OR, USA | |
| High Performance Liquid Chromatography System | ThermoFischer Scientific, Dreieich, GER | |
| ITC-16 | HEKA Elektronik, Lambrecht, GER | |
| KinematicaPolytron PT1300D homogenizer | ThermoFischer Scientific, Dreieich, GER | Catalogue #10764233 |
| Polychrome V | Till Photonics, Planegg-Martinsried, GER | |
| Proxeon nano-LC system | ThermoFischer Scientific, Dreieich, GER | |
| Reprosil 75 µm x 250 mm, 3 µm | Dr. Maisch GmbH, Ammerbuch, GER | |
| Spectrometer Thermo Orbitrap Fusion Mass | ThermoFischer Scientific, Dreieich, GER | |
| UVICO ultraviolett or visible light source | Rapp OptoElectronic, Hamburg, GER | |
| ***Software*** | | |
| ApE (A Plasmid Editor) | M. Wayne Davis, Utah, UT, USA | Version #10.55 |
| ImagePro Plus | MediaCybernetics, Roper, Sarasota, FL, USA | Version #6.0 |
| MaxQuant.Live | MaxQuant | Version #1.6.0.1 |
| Perseus | omicX | Version# 1.6.2.1 |
| Prism | GraphPad, San Diego, CA, USA | Version# 8 |
| Solis | Acal GmbH, Gröbenzell, GER | Version# 4.30.30034.0 |
| TIDA5.25 | HEKA Elektronik, Lambrecht, GER | Version# 5.25 |
| UGA-42 Firefly | Rapp OptoElectronic, Hamburg, GER | N/A |
